# Latitude Matters: A Global Phylogeographic Perspective on Climate-Driven Demographic Responses in Tarantulas

**DOI:** 10.1101/2025.06.08.658479

**Authors:** Aritra Biswas, Praveen Karanth

## Abstract

**Aim:** To investigate how past climate change has shaped the genetic diversity and demographic responses of tarantulas across latitudes, and to test whether climate–demography relationships vary with latitude.

**Location:** Global; spanning tropical to temperate regions.

**Taxon:** Tarantulas (family Theraphosidae).

**Methods:** We compiled mitochondrial *Cytochrome oxidase I* (COI) sequences for 48 tarantula species worldwide, including newly generated sequences, to estimate nucleotide diversity (π) and Tajima’s D. Species distribution models (SDMs) were constructed under present-day and Last Glacial Maximum (LGM) climatic conditions to quantify changes in habitat suitability since the LGM. Using generalized linear models (GLMs), we tested whether genetic and demographic metrics were associated with latitude and climate-driven habitat change, and whether their relationship varied with latitude.

**Results:** Pairwise correlations among latitude, habitat change, and genetic metrics showed no significant associations. However, GLMs revealed a significant interaction: the effect of habitat suitability change on Tajima’s D was strongly positive at high latitudes but negative or negligible at low latitudes. This indicates that demographic responses to past climate change varied latitudinally. Several high-latitude species showed genetic signatures of demographic expansion and range increase since the LGM.

**Main conclusions:** Our results support the hypothesis that species at higher latitudes experience stronger demographic fluctuations due to historical climate change, aligning with Darwin’s early predictions. Moreover, patterns of demographic growth in temperate taxa suggest that some species may benefit from recent warming, consistent with Janzen’s climatic variability hypothesis. These findings demonstrate that climate-driven genetic and demographic responses in tarantulas are shaped by latitude, highlighting the importance of integrating phylogeography with ecological niche modeling to understand species’ resilience under climate change.

## 1. Introduction

Understanding how climate change influences the distribution and diversity of life on Earth has become a central focus in contemporary ecological and evolutionary research. This growing interest stems from mounting empirical evidence that recent warming trends are driving widespread, and sometimes severe, biological responses (Cox et al., 2022; Malanoski et al., 2024; Mi et al., 2023; Michielsen et al., 2023; Veron et al., 2019). Yet, a critical question remains: are these effects uniform across the globe? Increasingly, studies suggest that species in different geographic regions, particularly across latitudinal gradients, respond differently to climate change, implying that its impact is spatially heterogeneous (Louthan et al., 2021; Mathes et al., 2024; Saupe et al., 2019).

This idea is rooted in a long-standing ecological debate. While Darwin posited that climate plays a stronger regulatory role in temperate regions (Darwin, 1859), Janzen argued that tropical species, with their narrower thermal tolerances, may be more vulnerable to even modest climatic shifts (Janzen, 1967). Though various theoretical frameworks have attempted to reconcile these views (Louthan et al., 2021; Zhang et al., 2022), large-scale empirical evaluations across broad taxonomic groups remain rare (Fonseca et al., 2023). As a result, there is a pressing need for macroecological and phylogeographic studies that explicitly test how species from different latitudes respond to environmental changes.

One of the most informative periods for understanding the long-term effects of climate change is the Quaternary, a geological period marked by repeated glacial–interglacial cycles (Williams et al., 1993). These cycles, particularly the Last Glacial Maximum (LGM, ~26,000–20,000 years ago), profoundly shaped Earth’s climates and ecosystems (Bennett, 1997; Williams et al., 1993). During the LGM, vast ice sheets covered much of the northern latitudes, global sea levels were lower, and many regions experienced arid conditions (Clark et al., 2009; Lambeck et al., 2014). Since then, Earth has undergone a dramatic warming trend, with average global surface temperatures rising by approximately 6°C—a rate of change that remains among the fastest in Earth’s history (Lambeck et al., 2014; Malanoski et al., 2024). This transition from the LGM to the present represents a natural experiment in rapid climate change, providing an ideal temporal framework to investigate its evolutionary and demographic consequences.

Genetic data offers a powerful lens through which to examine these impacts (G. M. Hewitt, 2004). DNA sequences preserve the signatures of past demographic events and have become essential tools for reconstructing historical population dynamics (G. M. Hewitt, 1993,1996, 1999, 2000, 2004; K Holder et al., 1999, 2000). Across diverse taxa, researchers have uncovered links between climate change since the LGM, population stability, and genetic diversity. Although the strength and direction of these relationships vary (Avise, 2000; Fonseca et al., 2023; G. M. Hewitt, 2004; Nogués-Bravo et al., 2010; Theodoridis et al., 2020).

Latitude is emerging as a key modulator of these effects. It shapes not only the magnitude and pace of climate change but also the spatial patterns of genetic diversity and population persistence (Avise, 2000; Bharti et al., 2023; Fonseca et al., 2023; French et al., 2023; Pearce-Higgins et al., 2015; Saupe et al., 2019). Generally, tropical regions exhibit greater climatic stability and less glacial disturbance than temperate and polar zones, which are subject to stronger seasonality, temperature fluctuations, and glacial advance and retreat (Barron, 1995; Dynesius & Jansson, 2000; Galván et al., 2025; Page & Shanker, 2020; Pianka, 1966). Together, these factors are likely to drive more pronounced shifts in the climatic niche space of species from higher latitudes than their tropical counterparts (Figure 1A, H1) (Barron, 1995; Dynesius & Jansson, 2000; Fonseca et al., 2023; Louthan et al., 2021; Page & Shanker, 2020; Pianka, 1966). Consequently, species in tropical zones are often thought to maintain more stable populations (Figure 1A, H2) (Barron, 1995; Fonseca et al., 2023; G. Hewitt, 2000; Louthan et al., 2021; Nogués-Bravo et al., 2010) and harbor higher intraspecific genetic diversity (Figure 1A, H3) (Avise, 2000; Fonseca et al., 2023; Hedrick & Kalinowski, 2000; G. M. Hewitt, 2004; Manel et al., 2020; Miraldo et al., 2016; Nogués-Bravo et al., 2010), while those at higher latitudes tend to experience more demographic fluctuations and reduced genetic diversity. Furthermore, climate-induced changes in a species’ ecological niche, termed climatic niche space, may negatively impact genetic diversity (Figure 1A, H4) (Chiocchio et al., 2024; de Pous et al., 2016; Fonseca et al., 2023; Hedrick & Kalinowski, 2000; G. Hewitt, 2000; Leigh et al., 2019; Lima et al., 2017) and destabilize populations (Figure 1A, H5) (Abreu-Jardim et al., 2021; Fonseca et al., 2023; G. M. Hewitt, 2004; Kanagaraj et al., 2023; Kardos et al., 2021; Leigh et al., 2019; Pearce-Higgins et al., 2015).

**Figure 1:**
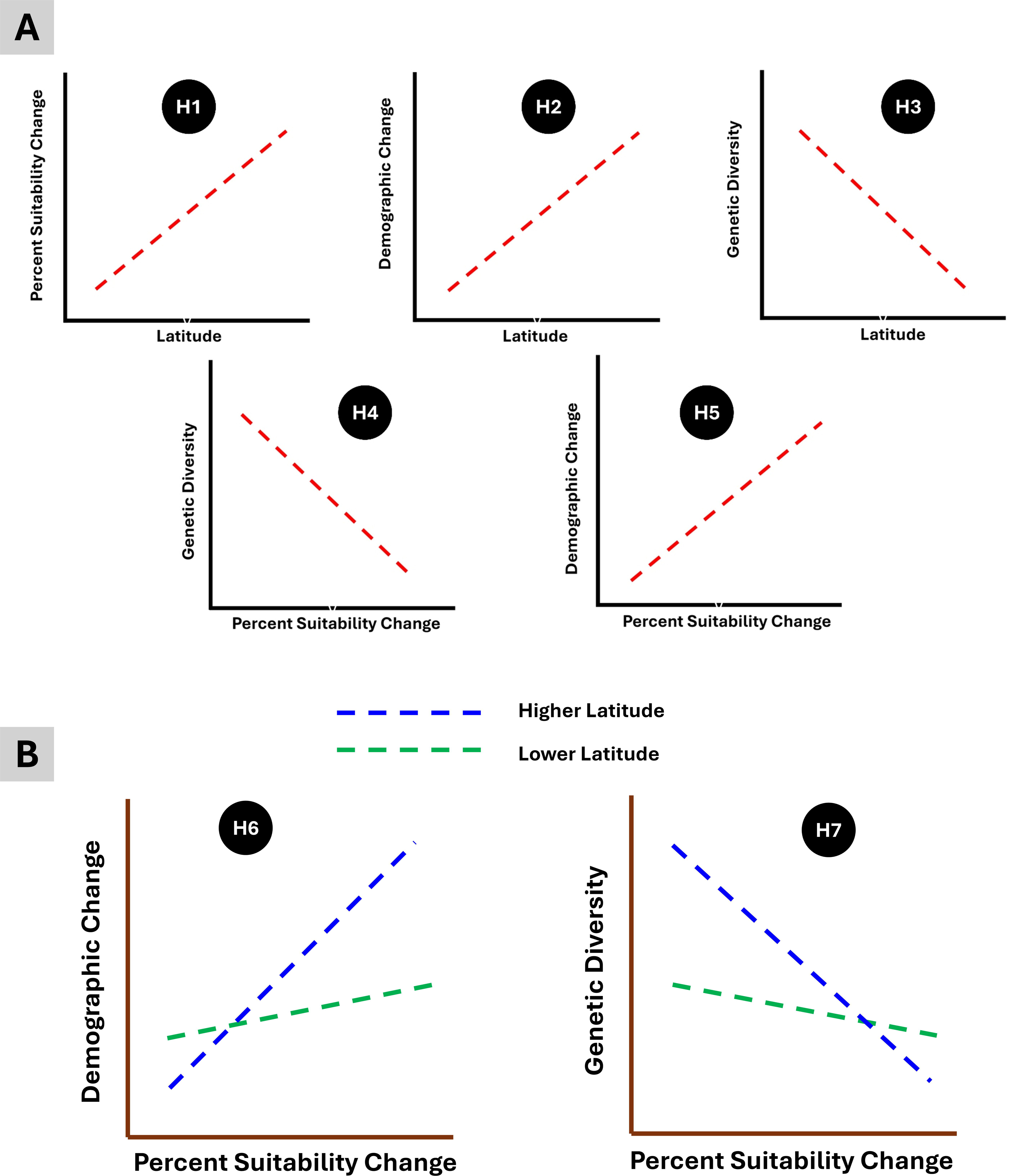
(A) The hypothesized relationships between all the variables. (B) The hypotheses regarding how the interaction of suitability change and latitude impact genetic diversity and demographic change.

Adding another layer of complexity, physiological sensitivity to climate change also appears to be latitude dependent, particularly for ectotherms (Deutsch et al., 2008; Louthan et al., 2021). Because these organisms rely entirely on external temperatures to regulate key physiological functions, they are especially vulnerable to warming climates (Cox et al., 2022; Deutsch et al., 2008; Mi et al., 2023). Studies suggest that tropical ectotherms often live near their upper thermal limits, rendering them particularly sensitive to even slight increases in temperature (Deutsch et al., 2008; Janzen, 1967). In contrast, temperate ectotherms typically possess broader thermal tolerances and often occupy environments cooler than their physiological optima, meaning that moderate warming could temporarily enhance their fitness (Deutsch et al., 2008; Janzen, 1967; Louthan et al., 2021). These differences in physiological vulnerability may translate into divergent demographic and genetic patterns across latitudes during periods of global warming. For example, via range expansion or increased population sizes in higher-latitude species, and contraction or decline in tropical taxa.

These patterns suggest that the demographic and genetic consequences of climate change are not only species-specific but also geographically patterned (Mathes et al., 2024; Pearce-Higgins et al., 2015). Synthesizing these insights, we hypothesize that the slope of the relationship between climatic niche change and demographic/genetic responses should differ across latitudes—being steeper in high-latitude taxa due to greater climatic volatility and glaciation history (Figure 1B, H6 and H7). However, these global hypotheses remain largely untested, as most studies investigating the effects of climate change on genetic and demographic patterns have been highly species-specific and geographically localized (Abreu-Jardim et al., 2021; de Pous et al., 2016; K Holder et al., 1999; Kanagaraj et al., 2023; Klimova & Landis, 2025; Lima et al., 2017; Page & Shanker, 2020; Sękiewicz et al., 2020).

Spiders of the family Theraphosidae (Infraorder: Mygalomorphae), commonly known as tarantulas, offer a compelling system in which to test these hypotheses. Comprising over 1000 species distributed across all continents except Antarctica, tarantulas are characterized by low dispersal abilities, high endemism, and traits like urticating hairs and venom (Biswas et al., 2023; Foley et al., 2021; Pocock, 1903). They face severe threats from habitat fragmentation and overexploitation for the pet trade, with many species currently listed as threatened by the IUCN (Branco & Cardoso, 2020; Rivera et al., 2024). So far, research on tarantulas has solely focused on taxonomy and systematics with some recent studies focusing on trait evolution and biogeographical aspects (Biswas et al., 2023; Biswas & Karanth, 2024, 2025; Foley et al., 2019, 2021). Despite their ecological and evolutionary significance, global assessments of how climate change has influenced their population history remain entirely absent.

In this study, we present the first global-scale analysis of the effects of post-LGM climate change on tarantula species. By integrating population genetic data, geographic distribution records, and species distribution models, we test a suite of well-defined hypotheses (Figure 1, H1–H7) linking climatic niche change, latitude, demographic history, and genetic diversity. Specifically, we explore both pairwise relationships between variables (Figure 1A, H1–H5) and multivariate models that account for interactive effects (Figure 1B, H6 and H7). Through this integrative framework, we aim to illuminate how past climate change has shaped the evolutionary and demographic trajectories of tarantulas and to contribute broader insights into the geography of genetic diversity and demographic responses in a rapidly changing world.

## 2. Materials and methods

### 2.1 Assembling DNA sequence data

We compiled mitochondrial cytochrome oxidase subunit I (COX I) sequences for tarantulas (family *Theraphosidae*) from GenBank, as this gene is the most commonly used molecular marker available for these spiders with well coverage across species. COX I has proven to be a reliable tool for phylogeographic studies in spiders (Derycke et al., 2010; Folmer et al., 1994; Fonseca et al., 2023; Hedin & Maddison, 2001), and it is widely used in population genetics to detect signals of genetic diversity and population divergence (Hajibabaei et al., 2007; Hu et al., 2019). To expand the global dataset, we generated additional COX I sequences for two Indian tarantula species—*Thrigmopoeus truculentus* and *Haploclastus kayi*. Field sampling was carried out in the Western Ghats, where spiders were collected and preserved in 70% ethanol. Genomic DNA was extracted from leg muscle tissue using the Qiagen DNeasy Blood and Tissue Kit (Qiagen, Hilden, Germany). A ~750 bp fragment of the COX I gene was amplified using the LCO and HCO primer pair, following the protocol outlined by Folmer et al. (1994). The newly obtained sequences were submitted to GenBank, and accession numbers for both the downloaded and newly generated sequences are provided in supplementary material 1. Overall, we assembled a total of 1208 sequences belonging to 48 tarantula species.

To reduce errors, we excluded species with fewer than five sequences from analyses. This threshold was informed by recent findings showing that the variance in nucleotide diversity and demographic estimates stabilizes beyond five samples per species (Barrow et al., 2021). Sequence alignments were performed separately for each species using the MUSCLE algorithm (Edgar, 2004) implemented in MEGA version 10.0 (Kumar et al., 2018). All alignments were translated into amino acid sequences to check for stop codons and ensure sequence quality before further analyses.

Figure 2A displays the global sampling locations, represented by the median latitude and longitude of each species with available genetic data.

**Figure 2:**
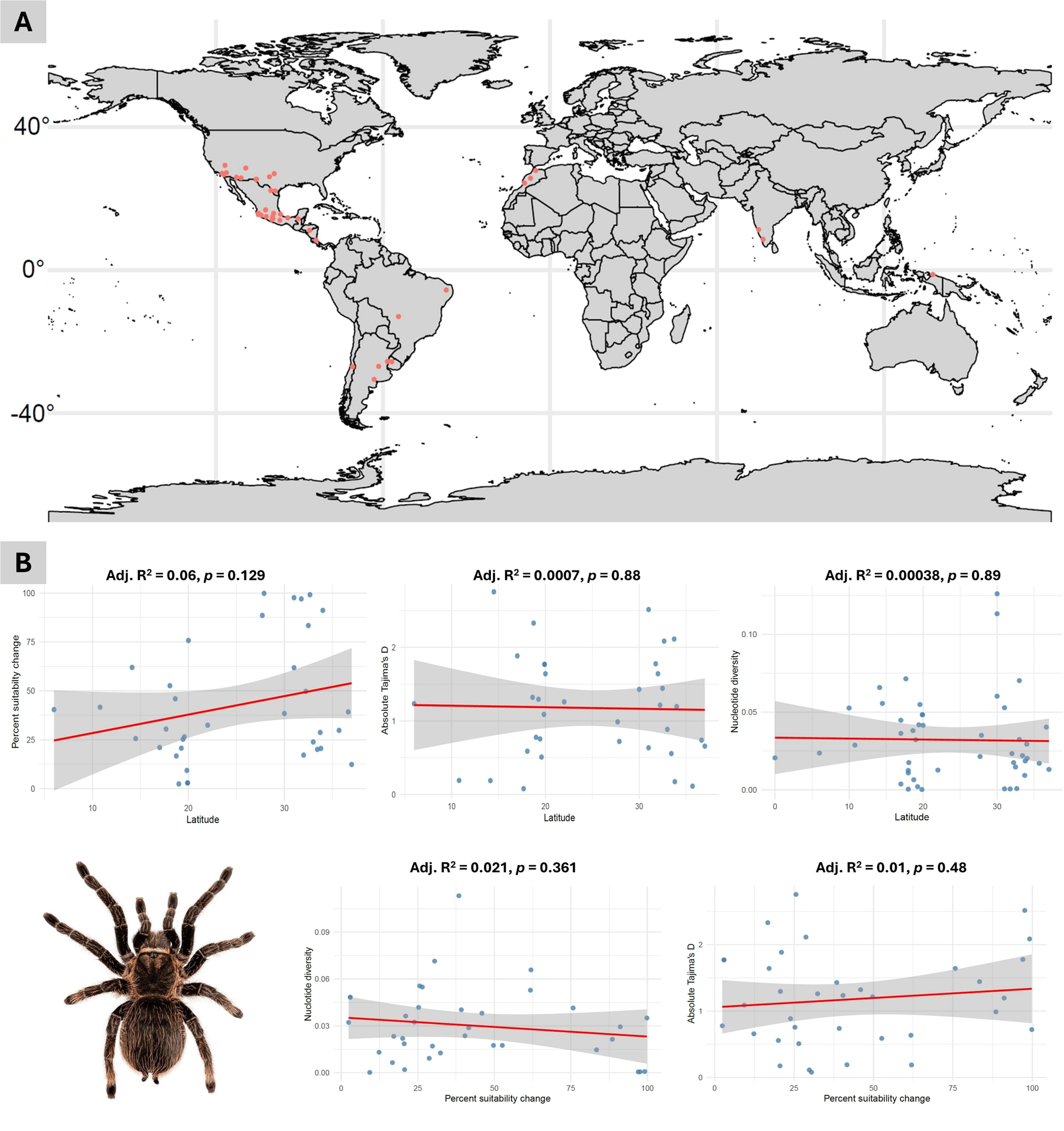
(A) Distribution of sampling locations (median latitudes and longitudes). (B) Pairwise correlation testing between all the variables

### 2.3 Calculating genetic diversity and assessing demographic change

To assess intraspecific genetic diversity, we calculated nucleotide diversity (π), which represents the average number of nucleotide differences per site between pairs of sequences (Nei & Li, 1979). While other measures such as the number of haplotypes or haplotype diversity are often used, they were not suitable for our dataset. This is because many sequences retrieved from GenBank are submitted as unique haplotypes per locality or vary in length, which limits the reliability of those metrics (Fonseca et al., 2023; Miraldo et al., 2016). Nucleotide diversity (π), in contrast, can be estimated reliably even from sequences of different lengths. We followed the approach described in Miraldo et al. (2016) and Fonseca et al. (2023), calculating pairwise diversity using the nuc.div function from the *pegas* R package (Paradis, 2010), and then averaged these values to obtain species-level estimates.

To detect signals of demographic change, we computed two neutrality statistics: Tajima’s D (Tajima, 1989) and R_2_ (Ramos-Onsins & Rozas, 2002). Tajima’s D compares two estimates of genetic diversity—mean pairwise differences (π) and the number of segregating sites (S). Under a stable population size, these values are expected to be similar. However, population expansions often lead to an excess of rare variants, resulting in lower π and more negative D values, whereas bottlenecks produce positive D values. Although Tajima’s D can also be influenced by selection, demographic processes are generally the primary interpretation in broad-scale phylogeographic analyses (Fonseca et al., 2023). We calculated Tajima’s D and associated p-values using the tajima.test function in *pegas* R package (Paradis, 2010).

R_2_ is another summary statistic sensitive to population expansion, based on the difference between the number of singletons and the average number of nucleotide differences. R_2_ values range from zero to one with values closer to zero indicating departures from neutrality due to expansion (Ramos-Onsins & Rozas, 2002). We computed R_2_ and its significance using the R2.test function in *pegas* (Paradis, 2010).

### 2.5 Species distribution models and projection to LGM

We compiled distribution data for all tarantula species with available genetic information from the Global Biodiversity Information Facility (GBIF; http://www.gbif.org/). To ensure data quality, we applied strict filtering procedures by removing duplicate records, missing values (NAs), and geographic points located in oceans. To mitigate potential spatial sampling bias, we employed spatial thinning by retaining only one occurrence point per 10 × 10 km grid cell. We then visually examined the cleaned distribution points by plotting them and comparing them to known species range maps from the IUCN Red List or published literature. Any points that lay substantially outside the known distribution ranges were manually removed. For downstream analyses, we retained only those species with ten or more occurrence records to ensure broad representation across genera while maintaining reasonably accurate estimates of species distributions (Carstens et al., 2018; van Proosdij et al., 2016).

From the final filtered occurrence data, we extracted 19 bioclimatic variables for each GPS point corresponding to both current climate conditions from WorldClim database (https://www.worldclim.org/) and the Last Glacial Maximum (LGM, ~22,000 years before present) from PaleoClim database (http://www.paleoclim.org/), all at a spatial resolution of 2.5 arc-minutes. To avoid multicollinearity, we excluded highly correlated variables (Pearson’s |r| ≥ 0.7), retaining only those variables that were uncorrelated for each species.

We used Maxent v3.4.4 (https://biodiversityinformatics.amnh.org/open_source/maxent/), a machine learning algorithm based on the principle of maximum entropy, to build species distribution models (SDMs). Maxent uses presence-only data and compares it to a random sample of background points to estimate the most uniform (i.e., least biased) distribution that still fits the observed occurrence data (Phillips et al., 2006; Phillips & Dudík, 2008). For each species, we randomly split the occurrence data into 70% training and 30% testing subsets, and used 10,000 randomly selected background points as pseudo-absences (Guillera-Arroita et al., 2014; Phillips et al., 2009). We performed five replicate model runs using cross-validation to assess consistency and applied a regularization multiplier of 1 to control model complexity. Model performance was evaluated using the area under the receiver operating characteristic curve (AUC). Only models with an AUC ≥ 0.70 were retained for further analysis, as models below this threshold were considered to have poor discriminatory power (Guillera-Arroita et al., 2014; Phillips et al., 2009, 2017).

The resulting SDMs were projected onto LGM climatic layers to estimate the distribution of suitable habitats during the glacial period. Model outputs were generated in cloglog format, which scales habitat suitability between 0 (lowest) and 1 (highest). These outputs were then processed to quantify changes in habitat suitability between the LGM and the present. We converted the continuous suitability maps into binary maps using two commonly used thresholding approaches: the “10 percentile training presence” (a) and “Equal training sensitivity and specificity” (b) (Phillips et al., 2006, 2009; Phillips & Dudík, 2008). For both thresholds, suitable habitat areas were calculated in square kilometers for the LGM and the present.

The percentage change in suitable habitat since the LGM was computed for each species using the formula:

{(absolute |Suitable_LGM − Suitable_Current|) / Suitable_LGM} × 100

This resulted in two estimates of climatic suitability change (percent_affected_area_a and percent_affected_area_b) corresponding to the two binarization methods.

All spatial data processing and post-processing steps were carried out using R statistical computing environment (R Core Team, 2021), with the following packages: sp (Pebesma & Bivand, 2005), dismo (Hijmans et al., 2024), raster (Hijmans, 2025), maptools (Bivand & Lewin-Koh, 2023), and rgdal (Bivand et al., 2023).

### 2.6 Statistical analysis

The final dataset comprised 48 tarantula species, including information on the number of DNA sequences, occurrence GPS points, median latitude of the distribution range, genetic diversity metrics (π, Tajima’s D, and R_2_), estimated suitable habitat areas under current and Last Glacial Maximum (LGM), and the percentage change in habitat suitability since the LGM calculated under two binarization thresholds. Of these 48 species, species distribution models (SDMs) could not be generated for 11 species due to insufficient occurrence data or poor model performance, leaving 37 species with usable SDM outputs. This curated dataset (Supplementary Material 2) was used for all downstream statistical analyses.

Before exploring relationships between variables, we first tested whether any of our key metrics were sensitive to sample size. We performed linear regressions to evaluate whether nucleotide diversity (π), Tajima’s D, and R_2_ were correlated with the number of sequences available per species. Similarly, we assessed whether the percentage change in habitat suitability since the LGM was correlated with the number of occurrence (GPS) points per species. These preliminary checks confirmed if our variables were significantly influenced by sample size, allowing us to proceed with further hypothesis testing.

To examine the ecological and evolutionary factors shaping genetic diversity, we used generalized linear models (GLM) to test a set of hypothesized relationships (Figure 1). First, we evaluated whether the percentage of habitat suitability changes since the LGM varied with latitude. Next, we modeled the effects of latitude, suitability change, and their interaction on genetic diversity indices (π, Tajima’s D, and R_2_). For each of these three genetic metrics, we fit three separate GLMs (Table 1): one including latitude as the sole predictor, one with suitability change as the sole predictor, and one with both predictors and their interaction. For ease of analysis and due to small sample sizes, we used the absolute values of both latitude and Tajima’s D.

**Table 1:**
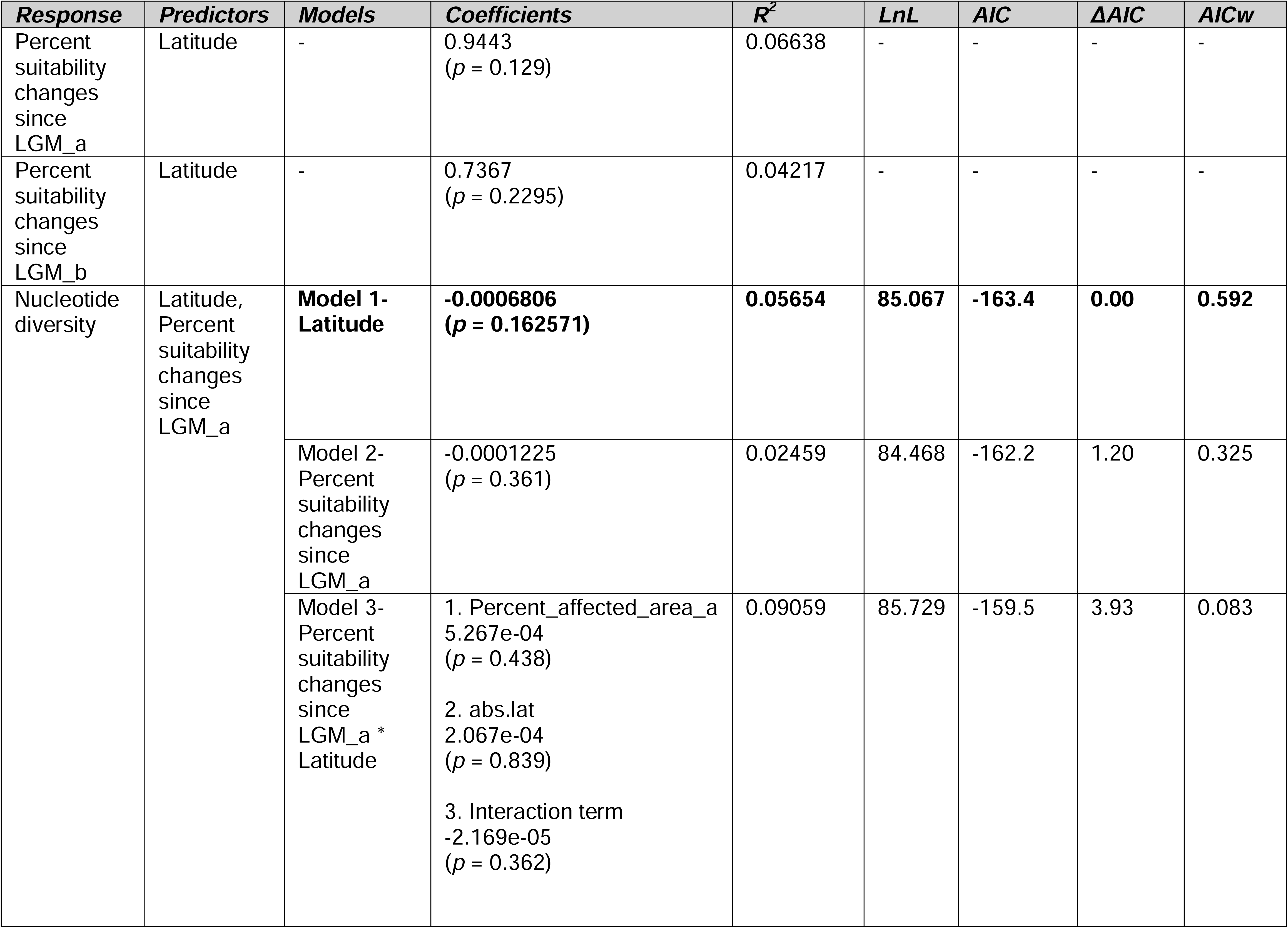
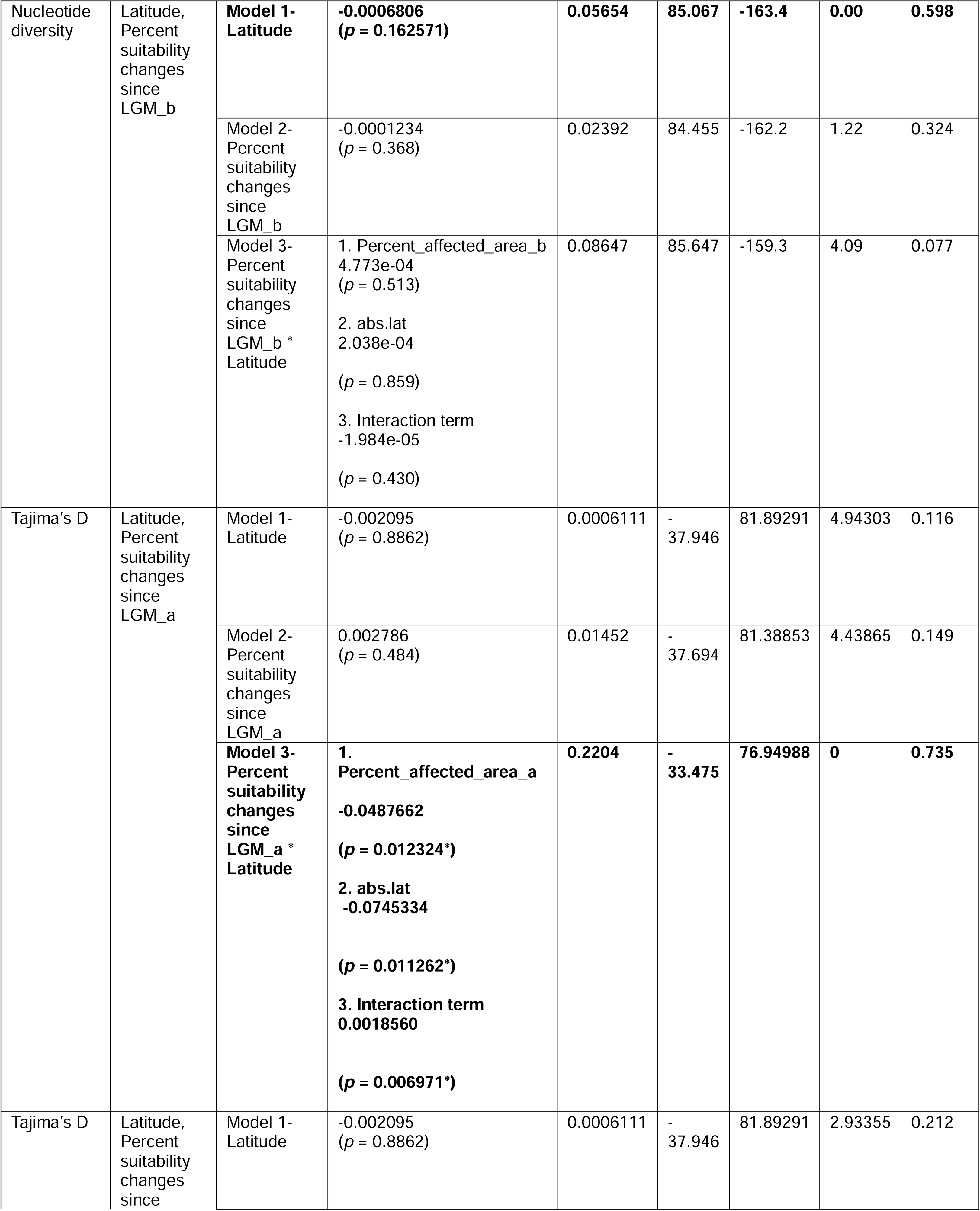
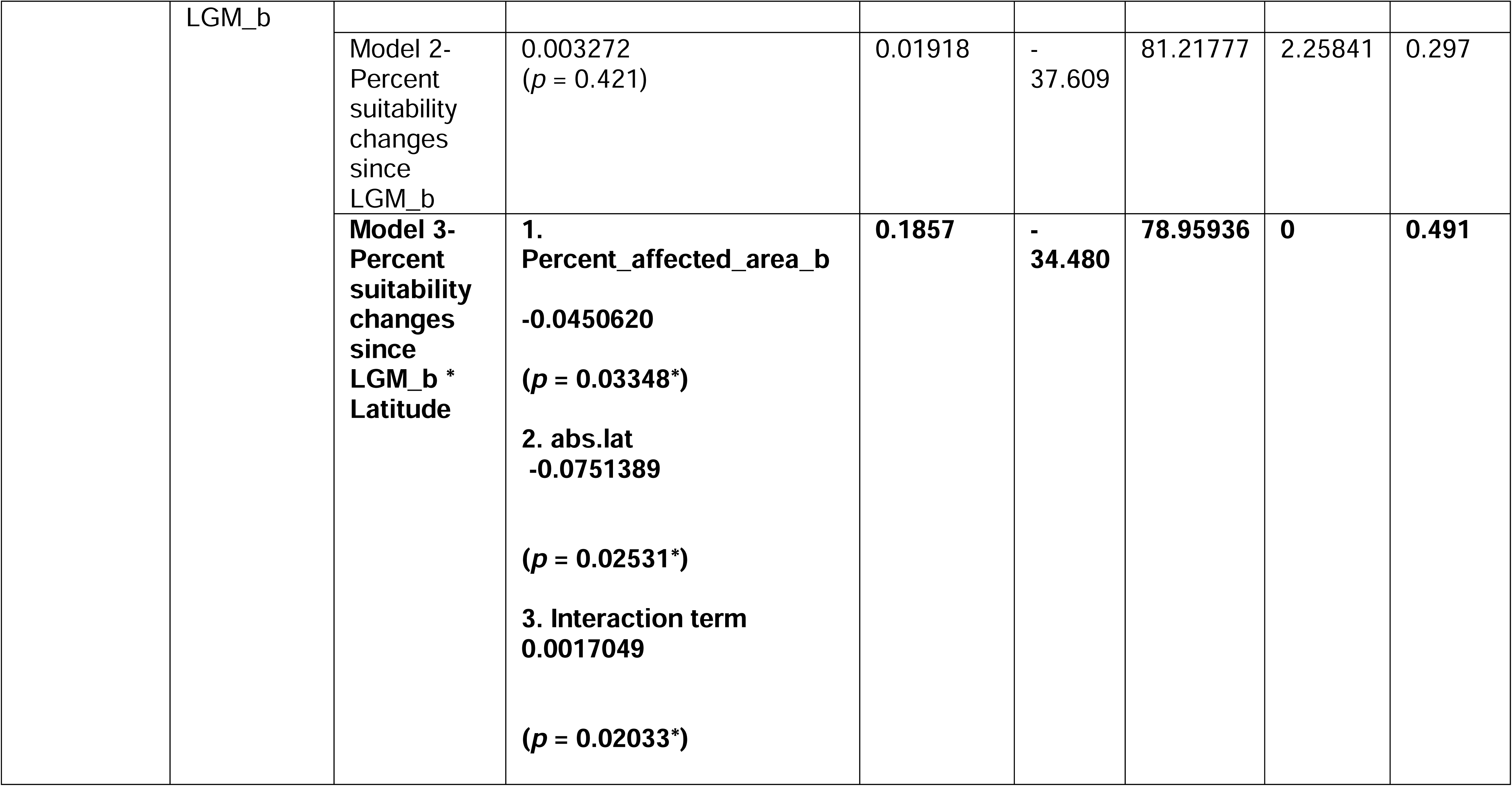
Results of the GLM analysis along with the coefficients estimated for every model, pseudo R^2^ value and AIC statistics. The best-fit models are highlighted in bold.

Model selection was based on Akaike Information Criterion (AIC), with lower AIC values indicating better model fit. For each response variable, we selected the best-fitting model based on AIC, and calculated pseudo-R^2^ values to assess the proportion of variance explained. Additionally, model coefficients were examined to evaluate the relative importance of predictors. All visualizations of model-predicted relationships were generated using the R package ggplot2 (Wickham, 2016).

## 3. Results

We first examined whether sample size influenced the genetic metrics used in this study. Correlation analysis revealed that nucleotide diversity (π) and Tajima’s D were not significantly affected by sample size (π: *r^2^* = 0.007, *p* = 0.55; Tajima’s D: *r^2^* = 0.06, *p* = 0.10). In contrast, the R_2_ statistic showed a strong negative correlation with sample size (*r^2^* = 0.45, *p* < 0.001), indicating sensitivity to the number of samples. Similarly, the percentage of climatically suitable area affected since the Last Glacial Maximum (LGM) was not associated with sample size (*r^2^* = 0.0009, *p* = 0.86). Given its sensitivity to sampling effort, we excluded the R_2_ metric from further analyses.

Next, we explored pairwise relationships among latitude, climatic suitability change, and genetic metrics. As hypothesized (Figure 1A, H1), we found a weak positive correlation between the percent change in suitable habitat since the LGM and latitude, although the relationship was not statistically significant (*r^2^* = 0.06, *p* = 0.129). Likewise, we found no significant relationship between latitude and either Tajima’s D (*r^2^* = 0.0007, *p* = 0.88) or nucleotide diversity (*r^2^* = 0.00038, *p* = 0.89). Percent suitability changes since the LGM also failed to predict Tajima’s D (*r^2^* = 0.021, *p* = 0.361) or nucleotide diversity (*r^2^* = 0.01, *p* = 0.48). Figure 2B presents the regression plots for these analyses.

We then used generalized linear models (GLMs) to further examine the relationships among these variables. Table 1 summarizes the model results, including coefficient estimates, significance levels, pseudo-*r^2^* values, and AIC-based model comparisons. For percent suitability change since the LGM, the GLM showed a positive association with latitude, but the relationship remained statistically non-significant, explaining only about 4–6% of the variation.

In the case of nucleotide diversity, the model including latitude alone (Model 1) performed better than models including suitability change alone (Model 2) or the interaction between latitude and suitability change (Model 3), under both binarization thresholds. Model 1 estimated a negative relationship between latitude and nucleotide diversity, but again, the effect was not statistically significant.

For Tajima’s D, however, the best-fitting model was Model 3, which included the interaction between latitude and percent suitability change. This interactive model explained a larger proportion of variance (*r^2^* = 22% and 18%) compared to Models 1 and 2 and outperformed them under both binarization thresholds. This result suggests that the effect of climatic suitability change on demographic history is latitude dependent.

Figure 3 visualizes the predicted relationships from Model 3 along with empirical data, using three latitude classes: low (<30%), mid (30–70%), and high (>70%). Panel A presents results for the 10-percentile training presence threshold, while Panel B shows results for the equal training sensitivity and specificity threshold. To check the robustness of these patterns, we also plotted Model 3 predictions using alternative latitude binning schemes (25–75% and 35–65% cut-offs), provided in Supplementary Material 3 (Figures S1 and S2).

**Figure 3:**
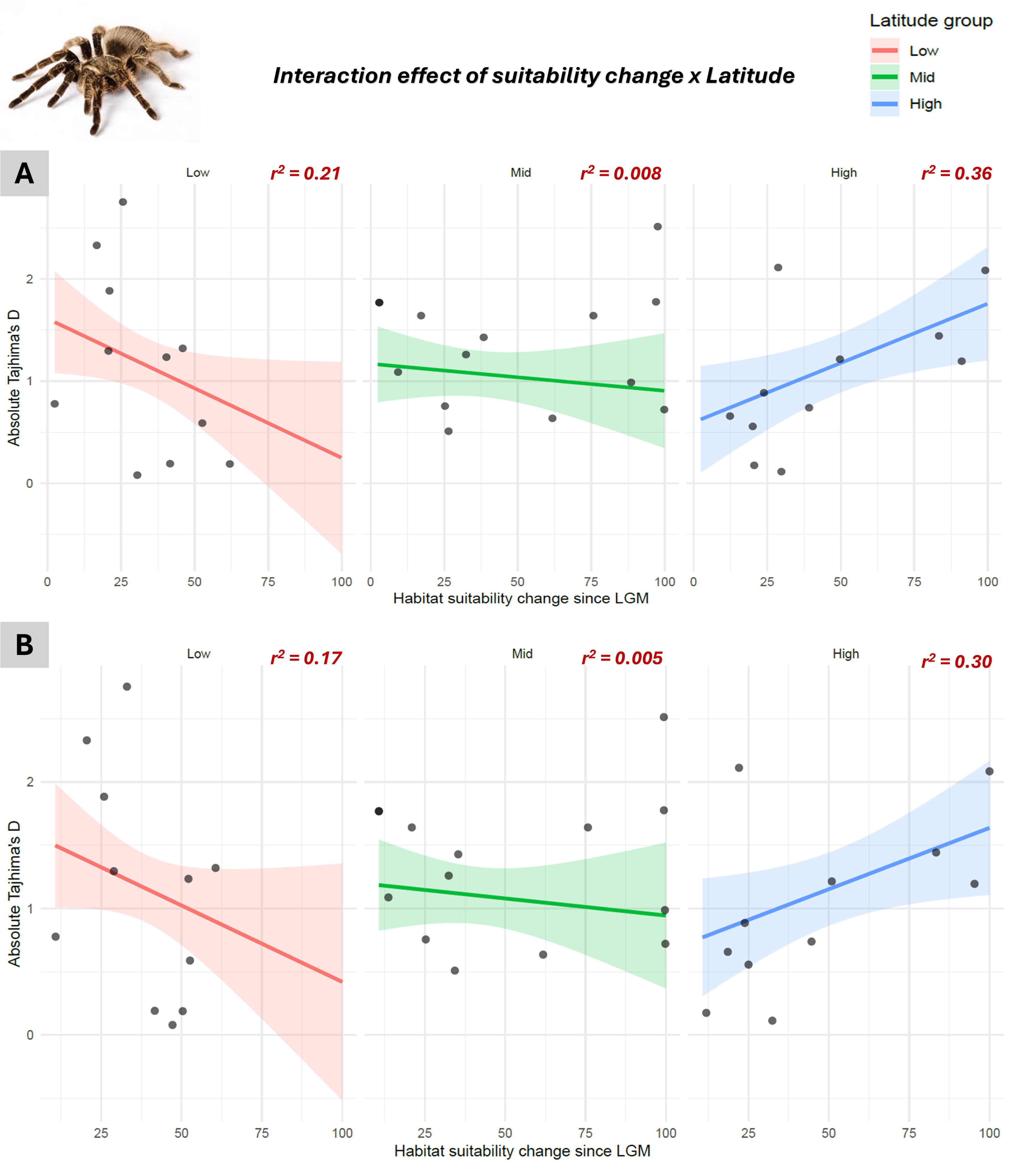
Visualization of the predicted effects of the habitat suitability change on Tajima’ D for different latitudes. Plot (A) depicts results for the 10 percentile training presence and (B) equal training sensitivity and specificity. The predicted values along with the 95% confidence interval for every latitude class are plotted along with the observed values to compare model fit. This plot is for the 30-70% latitude binning scheme where low latitude is <17[r], mid latitude is 17–33[r] and high latitude is >33[r].

Finally, we ranked species based on their nucleotide diversity values and demographic responses. Figure 4A shows the species ranking by nucleotide diversity, indicating whether each species experienced a contraction or expansion in suitable habitat according to the niche models. For a few species, the direction of change could not be determined due to conflicting results from different binarization thresholds. Figure 4B ranks species by Tajima’s D values, highlighting those that likely underwent population expansion or contraction. Figure 4C presents species ranked by the extent of climatically suitable area affected since the LGM.

**Figure 4:**
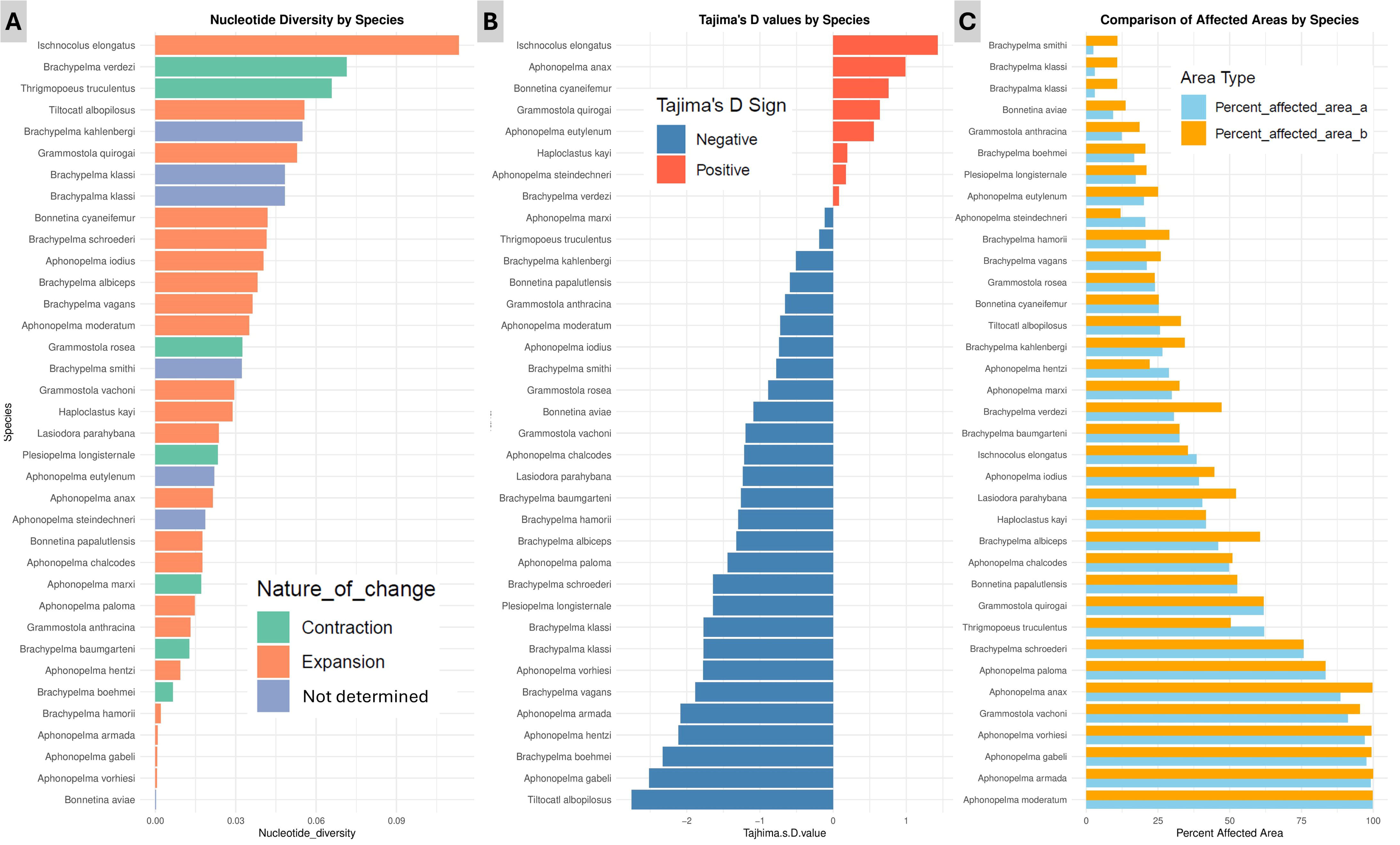
Ranking of species according to (A) nucleotide diversity, (B) Tajima’s D value and (C) percentage of habitable area affected since LGM according to two different binarization cut-offs.

Supplementary Material 4 presents a comparison of the raw Maxent outputs depicting climatic suitability during the Last Glacial Maximum (LGM) and the present day for all species. The scripts used for spatial analysis and GLM are provided in the supplementary material 5.

## 4. Discussion

### 4.1 Climate change impacts are shaped by latitude

Our global analysis of tarantulas offers empirical support for the idea that climate-driven demographic responses are latitude dependent, even if simple pairwise relationships failed to reveal significant patterns. This initial lack of correlation between latitude, climatic suitability change, and genetic metrics such as nucleotide diversity and Tajima’s D was unexpected, especially given previous studies across diverse taxa including plants, vertebrates, and insects—that have consistently documented latitudinal gradients in intraspecific genetic diversity and climate-linked demographic shifts (Avise, 2000; Fonseca et al., 2023; Nogués-Bravo et al., 2010). In our case, however, it was only through a multivariate framework that the geographic signature of climate change on tarantula demography became evident.

The interaction model (Model 3) between latitude and percent suitability change provided the best fit for explaining Tajima’s D, revealing a nuanced pattern: at low latitudes, increased climatic instability appears to drive less Tajima’s D values (indicative of population consistency), while at high latitudes, the effect is reversed as greater suitability change is associated with higher Tajima’s D values, implying population fluctuations. This latitude-specific modulation of demographic responses aligns with Hypothesis H6 (Figure 1B), which predicted that the slope of the relationship between climate change and demography varies across latitudes.

Importantly, this pattern remained robust across different latitude binning schemes (25–75%, 30–70%, and 35–65%), reinforcing the consistency of the demographic signal, especially at higher latitudes. While the strength of the negative relationship between suitability changes and Tajima’s D at low latitudes weakened under some binning thresholds, the positive association at higher latitudes remained stable (Figure 3; Supplementary Figures S1–S2). This provides compelling support for the idea that species in temperate zones, shaped by a history of glaciations and stronger seasonal variation, are more demographically sensitive to post-glacial climatic shifts.

Our results are consistent with the predictions of both Darwin and Janzen. Darwin emphasized the role of environmental variability in temperate zones in regulating populations (Darwin, 1859), a notion echoed here by the stronger demographic instability observed in high-latitude tarantulas. Meanwhile, Janzen’s “mountain passes are higher in the tropics” hypothesis highlights how narrow thermal tolerances in tropical species render them more vulnerable to climatic shifts (Janzen, 1967). Although we did not find strong evidence of demographic contraction in tropical species, our results suggest limited demographic fluctuation in the tropics—possibly reflecting the relative climatic stability of these regions since the Last Glacial Maximum.

Interestingly, many temperate tarantula species like *Aphonopelma*, *Brachypelma*, and *Grammostola* spp. showed negative Tajima’s D values alongside evidence of range expansion in our species distribution models (Figures 4B–C; Supplementary Material 4). This pattern is consistent with Janzen’s prediction that warming may allow temperate ectotherms to expand their ranges due to broader thermal tolerance and historical occupancy of suboptimal thermal environments (Deutsch et al., 2008; Louthan et al., 2021).

### 4.2 Limited explanatory power, hidden factors, and data gaps

While our best-fit model explained up to 18–22% of the variance in Tajima’s D, a large portion of demographic variation remains unexplained. This modest explanatory power could reflect the influence of additional, unmeasured factors including local-scale ecological conditions, species-specific life-history traits, microhabitat preferences, or historical biogeographic events unrelated to climate. Furthermore, mitochondrial DNA alone may not capture the full spectrum of population genetic responses, especially given its maternal inheritance and susceptibility to lineage sorting and selective sweeps (Guo et al., 2020; McGuire et al., 2007). Incorporating genome-wide data in future studies could offer more comprehensive insights.

Our study also underscores a notable sampling bias that limits generalization: as shown in Figure 2A, most sequencing efforts to date have been concentrated in the New World (especially the Americas), while the Old World remains critically undersampled. This introduces a longitudinal bias in our dataset that could not be avoided, but it simultaneously highlights a critical knowledge gap. There is a severe lack of DNA data from Old World tarantula species, particularly from Africa, Asia, and Australasia. This prevents any robust phylogeographic or demographic inferences from being drawn for these regions. Generating more primary sequence data from these underrepresented areas is therefore essential for a comprehensive understanding of how global climate change has influenced tarantula diversity and demography.

Expanding both the taxonomic and geographic breadth of sampling will be vital in future efforts to refine the ecological and evolutionary conclusions of this study. Broader data coverage could help clarify whether the patterns observed here such as latitude-dependent demographic responses are indeed global features or artifacts of biased sampling.

### 4.3 Implications for tarantula conservation

Our findings have direct implications for tarantula conservation. Despite being globally distributed, tarantulas are characterized by low dispersal abilities, high endemism, and narrow habitat requirements (Biswas & Karanth, 2024; Foley et al., 2021; Pocock, 1903; Rivera et al., 2024)—traits that make them particularly vulnerable to climate-induced habitat shifts. The fact that many high-latitude species appear to be expanding their ranges suggests a potential for demographic recovery under warming scenarios. However, this same warming may destabilize populations in more climatically stable tropical zones by subtly eroding suitable microhabitats or amplifying the effects of habitat fragmentation.

Conservation strategies should therefore account for the geographic and climatic context of each species. Prioritizing habitat corridors in tropical regions may help maintain genetic connectivity and buffer against future climatic volatility. Simultaneously, monitoring demographic trajectories of expanding temperate species can help assess long-term viability, especially in landscapes undergoing rapid land-use change.

## 5. Conclusion

This study presents the first global phylogeographic assessment of climate-driven demographic responses in tarantulas, a diverse yet understudied group of low-dispersal, ecologically specialized spiders. By integrating genetic data, species distribution modeling, and climatic niche dynamics across latitudes, we show that demographic responses to past climate change are geographically structured. Specifically, the relationship between climatic suitability changes and demographic history, as reflected by Tajima’s D, varies with latitude—being strongest in high-latitude species and relatively weak in low latitude taxa. These findings support long-standing ecological theories proposed by Darwin and Janzen.

Our results offer broader insights into how ectotherms with limited dispersal may respond to future climate change and underscore the critical importance of geographic context in demographic outcomes. We also highlight a major gap in biodiversity data, especially from the Old World, emphasizing the need for expanded sampling and sequencing. Overall, this work provides a framework for integrating macroecological theory with conservation and calls for regionally tailored strategies to protect vulnerable species in a warming world.

## Acknowledgements

We are grateful to the Forest Departments of Karnataka (C1(D)/WL/CR-30/2010-11), Kerala (WL 12-5806/2010), and Tamil Nadu (WL 5/42749/2010) for granting permission to collect specimens. We also thank Amatya Sharma, David Raju, Sajith Palat, Avrajjal Ghosh, R. Chaitanya, Vivek Cyriac and Madhura Agashe for their assistance with field sampling. AB wants to thank the International Society of Arachnology (ISA) and American Arachnological Society (AAS) for providing student grants.

## Conflict of interest

The authors declare that no conflict of interest exists.

## Author contributions

AB and PK conceived the study. AB conducted sampling, generated data, performed analysis and wrote the first draft. PK vetted and modified the manuscript.

## Data availability statement

All the newly generated sequences in this study were submitted to GenBank under the accession numbers PV630228–PV630258 and PV628768–PV628771.

## Notes

### Competing Interest Statement

The authors have declared no competing interest.

