## Supplementary material 1 for "Latitude Matters: A Global Phylogeographic Perspective on Climate-Driven Demographic Responses in Tarantulas"

| **Species** | **Cox I accession numbers** |
| --- | --- |
| *Aphonopelma anax* | JF803303.1  JF803304.1  JF803305.1  JF803306.1  JF803307.1  JF803308.1  JF803309.1  JF803310.1  JF803311.1  JF803312.1  JF803313.1  JF803314.1  JF803315.1  JF803316.1  JF803317.1  JF803318.1  JF803319.1  JF803320.1  JF803321.1  JF803322.1  JF803323.1  JF803324.1  JF803325.1  JF803326.1  JF803327.1  KC664899.1  KC664900.1  KC664901.1  KC664931.1  KC664932.1  KC664933.1  KC664972.1  KC664973.1  KC665125.1  KC665126.1  KC665127.1  KC665128.1  KC665129.1  KC665130.1  KC665131.1  KC665133.1  KC665134.1  KC665135.1  KC665136.1  KC665169.1  KC665170.1  KC665171.1  KC665172.1  KC665173.1  KC665174.1  KC665201.1  KC665202.1  KC665203.1  KC665204.1  KC665205.1  KC665206.1  KC665227.1  KC665250.1  KC665268.1 |
| *Aphonopelma armada* | JF803370.1  JF803371.1  JF803372.1  JF803373.1  JF803374.1  JF803375.1  JF803376.1  JF803377.1  JF803378.1  JF803379.1  JF803380.1  JF803381.1  JF803382.1  JF803383.1  JF803384.1  JF803385.1  JF803386.1  KC664790.1  KC664948.1  KC664949.1  KC664950.1  KC664951.1  KC664952.1  KC664953.1  KC664954.1  KC664955.1  KC664979.1  KC664980.1  KC665132.1  KC665153.1  KC665154.1  KC665155.1  KC665156.1  KC665157.1  KC665158.1  KC665159.1  KC665160.1  KC665161.1  KC665162.1  KC665168.1  KC665225.1  KC665226.1  KC665247.1  KC665248.1  KC665253.1  KC665256.1  KC665265.1  KC665266.1  KC665267.1  KC665276.1  KC665344.1  KC665347.1  KC665351.1  KC665352.1  KC665353.1  KC665354.1  KC665356.1  KC665357.1  KC665359.1  KC665360.1  KC665362.1  KC665363.1  KC665364.1 |
| *Aphonopelma chalcodes* | KC664794.1  KC664813.1  KC664835.1  KC664853.1  KC664854.1  KC664863.1  KC664875.1  KC664877.1  KC664882.1  KC664892.1  KC664919.1  KC664920.1  KC664922.1  KC664923.1  KC664924.1  KC664925.1  KC664981.1  KC664982.1  KC664983.1  KC664984.1  KC664985.1  KC664986.1  KC664987.1  KC664988.1  KC664989.1  KC664990.1  KC664991.1  KC664992.1  KC664993.1  KC664994.1  KC664995.1  KC664996.1  KC664997.1  KC664998.1  KC664999.1  KC665001.1  KC665002.1  KC665003.1  KC665046.1  KC665047.1  KC665048.1  KC665049.1  KC665050.1  KC665051.1  KC665052.1  KC665053.1  KC665054.1  KC665055.1  KC665056.1  KC665057.1  KC665069.1  KC665070.1  KC665073.1  KC665074.1  KC665075.1  KC665076.1  KC665089.1  KC665090.1  KC665091.1  KC665092.1  KC665194.1  KC665257.1 |
| *Aphonopelma eutylenum* | JF803417.1  KC664845.1  KC664861.1  KC664880.1  KC665199.1  KC665280.1  KC665281.1  KC665282.1  KC665283.1  KC665306.1  KC665307.1  KC665311.1  KC665312.1  KC665313.1  KC665314.1  KC665315.1  KC665316.1  KC665317.1  KC665318.1  KC665320.1  KC665323.1  KC665324.1  KC665326.1  KC665330.1  KC665332.1  KC665333.1  KC665335.1  KC665336.1  KC665337.1  KC665340.1  KC665380.1  KC665381.1  KC665382.1  KC665383.1  KC665384.1  KC665385.1  KC664840.1 |
| *Aphonopelma gabeli* | KC664798.1  KC664805.1  KC664810.1  KC664829.1  KC664865.1  KC664890.1  KC664939.1  KC664942.1  KC664943.1  KC664944.1  KC664946.1  KC665007.1  KC665008.1  KC665009.1  KC665010.1  KC665011.1  KC665012.1  KC665013.1  KC665014.1  KC665017.1  KC665018.1  KC665019.1  KC665020.1  KC665022.1  KC665023.1  KC665024.1  KC665025.1  KC665026.1  KC665027.1  KC665028.1  KC665030.1  KC665036.1  KC665037.1  KC665038.1  KC665039.1  KC665041.1  KC665042.1  KC665043.1  KC665045.1  KC665058.1  KC665059.1  KC665061.1  KC665062.1  KC665063.1  KC665064.1  KC665065.1  KC665066.1  KC665067.1  KC665068.1  KC665071.1  KC665072.1  KC665163.1  KC665164.1  KC665165.1  KC665166.1  KC665167.1  KC665195.1  KC665243.1  KC665246.1  KC665249.1  KC665345.1  KC665346.1  KC665348.1  KC665349.1  KC665355.1  KC665350.1  KC665395.1  KC665396.1  KC665397.1  KC665398.1  KC665399.1  KC665401.1  KC664782.1 |
| *Aphonopelma hentzi* | KC664941.1 KC664957.1 KC664971.1 KC665461.1 KC665361.1 KC665275.1 KC665274.1 KC665272.1 KC665264.1 KC665263.1 KC665262.1 KC665261.1 KC665260.1 KC665254.1 KC665252.1 KC665242.1 KC665236.1 KC665230.1 KC665224.1 KC665223.1 KC665222.1 KC665221.1 KC665220.1 |
| *Aphonopelma iodius* | JX244811.1 KC665460.1 KC665458.1 KC665453.1 KC665452.1 KC665451.1 KC665450.1 KC665449.1 KC665448.1 KC665447.1 KC665372.1 KC665371.1 KC665370.1 KC665300.1 KC665299.1 KC665298.1 KC665297.1 KC665296.1 KC665118.1 KC665117.1 KC664928.1 KC664908.1 KC664784.1 |
| *Aphonopelma joshua* | JX946016.1 JX946017.1 JX946018.1 JX946019.1 KP677300.1 KP677301.1 KC664913.1 KC664914.1 KC665304.1 KC665403.1 |
| *Aphonopelma marxi* | KC664788.1 KC664789.1 KC664806.1 KC664837.1 KC664852.1 KC664873.1 KC664897.1 KC664926.1 KC664927.1 KC665100.1 KC665105.1 KC665106.1 KC665107.1 KC665108.1 KC665109.1 KC665110.1 KC665111.1 KC665112.1 KC665116.1 KC665187.1 JF803328.1 KC664784.1 JX244811.1 KC665460.1 KC665458.1 KC665453.1 KC665452.1 KC665451.1 KC665450.1 KC665449.1 KC665448.1 KC665447.1 KC665372.1 KC665371.1 KC665370.1 KC665300.1 KC665299.1 KC665298.1 KC665297.1 KC665296.1 KC665118.1 KC665117.1 KC664928.1 KC664908.1 JX946016.1 JX946017.1 JX946018.1 JX946019.1 KP677300.1 KP677301.1 KC664913.1 KC664914.1 KC665304.1 KC665403.1 KC665418.1 |
| *Aphonopelma moderatum* | JF803409.1 JF803410.1 JF803408.1 KC664807.1 KC664826.1 KC664843.1 KC664902.1 KC664903.1 KC664904.1 KC664905.1 KC664907.1 KC664935.1 KC664936.1 KC665186.1 KC665197.1 KC665233.1 JF803328.1 KC664784.1 JX244811.1 KC665460.1 KC665458.1 KC665453.1 KC665452.1 KC665451.1 KC665450.1 KC665449.1 KC665448.1 KC665447.1 KC665372.1 KC665371.1 KC665370.1 KC665300.1 KC665299.1 KC665298.1 KC665297.1 KC665296.1 KC665118.1 KC665117.1 KC664928.1 KC664908.1 JX946016.1 JX946017.1 JX946018.1 JX946019.1 KP677300.1 KP677301.1 KC664913.1 KC664914.1 KC665304.1 KC665403.1 KC665418.1 KC664788.1 |
| *Aphonopelma paloma* | JX946014.1 JX946015.1 KC665424.1 KC665425.1 JX946013.1 JF803328.1 KC665460.1 JX244811.1 KC665403.1 JX946016.1 KC665418.1 KC665425.1 KC664788.1 JF803409.1 KC665233.1 |
| *Aphonopelma steindechneri* | KC664791.1 KC664792.1 KC664817.1 KC664819.1 KC664841.1 KC664842.1 KC664856.1 KC664888.1 KC664894.1 KC665284.1 KC665288.1 KC665289.1 KC665290.1 KC665291.1 KC665292.1 KC665308.1 KC665309.1 KC665310.1 JF803328.1 JX244811.1 KC665460.1 KC664784.1 KC665403.1 JX946016.1 KC665418.1 KC664788.1 JF803409.1 KC665233.1 |
| *Aphonopelma vorhiesi* | KC664781.1 KC664855.1 KC664859.1 KC664860.1 KC664866.1 KC664867.1 KC664870.1 KC664871.1 KC664872.1 KC664891.1 KC664893.1 KC664895.1 KC665005.1 KC665006.1 KC665015.1 KC665016.1 KC665031.1 KC665033.1 KC665086.1 KC665087.1 JF803328.1 JX244811.1 KC665460.1 KC664784.1 JX946016.1 KC665403.1 KC664788.1 JF803409.1 KC665233.1 KC664791.1 KC664792.1 KC664817.1 KC664819.1 KC664841.1 KC664842.1 KC664856.1 KC664888.1 KC664894.1 KC665284.1 KC665288.1 KC665289.1 KC665290.1 KC665291.1 KC665292.1 KC665308.1 KC665309.1 KC665310.1 KC665418.1 KC665338.1 KY017963.1 |
| *Bonnetina alagoni* | KU664207.1 KU664208.1 KU664209.1 KU664210.1 KU664211.1 JF803328.1 JX244811.1 KC665460.1 KC664784.1 JX946016.1 KC665403.1 KC664788.1 JF803409.1 KC665233.1 KC664791.1 KC664855.1 KC664859.1 KC664860.1 KC664866.1 KC664867.1 KC664870.1 KC664871.1 KC664872.1 KC664891.1 KC664893.1 KC664895.1 KC665005.1 KC665006.1 KC665015.1 KC665016.1 KC665031.1 KC665033.1 KC665086.1 KC665087.1 KC665418.1 KC665338.1 KY017963.1 |
| *Bonnetina aviae* | KP757184.1 KP757185.1 KP757220.1 KP757221.1 KP757222.1 KP757223.1 KP757224.1 KP757225.1 KU664204.1 JF803328.1 JX244811.1 KC665460.1 KC664784.1 JX946016.1 KC665403.1 KC664788.1 JF803409.1 KC665233.1 KC664791.1 KC664855.1 KC664859.1 KC664860.1 KC664866.1 KC664867.1 KC664870.1 KC664871.1 KC664872.1 KC664891.1 KC664893.1 KC664895.1 KC665005.1 KC665006.1 KC665015.1 KC665016.1 KC665031.1 KC665033.1 KC665086.1 KC665087.1 KC665418.1 KC665338.1 KY017963.1 KU664207.1 KU664208.1 KU664209.1 KU664210.1 KU664211.1 |
| *Bonnetina cyaneifemur* | KP757190.1 KP757191.1 KP757192.1 KP757193.1 KP757231.1 KP757232.1 KP757233.1 |
| *Bonnetina minax* | KP757195.1 KP757196.1 KP757236.1 KP757237.1 KP757238.1 |
| *Bonnetina papalutlensis* | KP757197.1 KP757198.1 KP757199.1 KP757200.1 KP757201.1 KP757202.1 KP757203.1 KP757204.1 KP757205.1 KP757206.1 KP757207.1 KP757208.1 KP757209.1 KP757210.1 KP757211.1 KP757212.1 KP757239.1 KP757240.1 KP757241.1 KP757242.1 KP757243.1 KP757244.1 KP757245.1 |
| *Bonnetina tanzeri* | KP757213.1 KP757214.1 KP757215.1 KP757216.1 KP757217.1 KP757246.1 KP757247.1 KP757248.1 KP757249.1 KP757250.1 KP757251.1 KP757252.1 KP757253.1 KP757254.1 KU664216.1 |
| *Bonnetina tindoo* | KP757188.1 KP757226.1 KP757227.1 KP757228.1 KP757229.1 KP757213.1 KP757214.1 KP757215.1 KP757216.1 KP757217.1 KP757246.1 KP757247.1 KP757248.1 KP757249.1 KP757250.1 KP757251.1 KP757252.1 KP757253.1 KP757254.1 KU664216.1 |
| *Bonnetina vittata* | KP757218.1 KP757255.1 KP757256.1 KU664212.1  KX835171.1 |
| *Brachypalma klassi* | KT995340.1 KT995329.1 KT995346.1 KT995330.1 KT995349.1 KT995377.1 KT995393.1 DQ224251.1 DQ224250.1 |
| *Brachypelma albiceps* | OR687434.1 KT995354.1 KT995391.1 KT995398.1 KT995331.1 KT995384.1 |
| *Brachypelma baumgarteni* | KT995386.1 KT995395.1 KT995402.1 KT995382.1 KT995332.2 DQ224253.1 |
| *Brachypelma boehmei* | KT995337.1 MH362809.1 KT995343.1 KT995359.1 DQ224246.1 DQ224247.1 |
| *Brachypelma hamorii* | KT995401.1  KT995378.1  KT995334.1  KT995325.1  KT995381.1  KT995387.1 |
| *Brachypelma kahlenbergi* | KT995356.1  KT995361.1  KT995372.1  KT995396.1  KT995339.1  KT995347.1  KT995362.1  KT995388.1  KT995335.2  KT995366.1 |
| *Brachypelma schroederi* | KT995399.1  KT995333.1  KT995345.1  KT995376.1  KT995390.1  KT995394.1  KT995370.2 |
| *Brachypelma smithi* | DQ224252.1  KT995374.2  KT995363.2  KT995400.1  KT995375.1  KT995364.1  KT995385.1  KT995380.1 |
| *Brachypelma vagans* | MK270584.1  MK270585.1  MK270586.1  MK270587.1  AJ584615.1  AJ584616.1  AJ584617.1  AJ584618.1  AJ584620.1  AJ584621.1  AJ584624.1  AJ584625.1  AJ584626.1  AJ584628.1  AJ584629.1  AJ584630.1  AJ584631.1  AJ584632.1  AJ584633.1  AJ584635.1  AJ584636.1  AJ584627.1  AJ584634.1  AJ584623.1  AJ584619.1  AJ584622.1  KT995327.1  KT995341.1  KT995355.1  KT995358.1  KT995397.1  KT995342.1  KT995336.1 |
| *Brachypelma verdezi* | KT995351.1  KT995369.1  KT995328.1  KT995367.1  KT995379.1  KT995383.1  KT995389.1  KT995392.1  KT995352.2  KT995344.2  KT995368.1 |
| *Grammostola anthracina* | KT965202.1  KT965236.1  KT965237.1  KT965238.1  KT965245.1  KT965247.1  KT965250.1  KT965255.1  KT965258.1  KT965263.1  KT965248.1  KT965246.1  KT965232.1  KT965226.1  KT965227.1  KT965228.1  KT965225.1  KT965267.1  KT965219.1  KT965222.1  KT965223.1  KT965224.1 |
| *Grammostola pulchra* | KT965206.1  KT965207.1  KT965209.1  KT965210.1  KT965211.1  OM670234.1  OM670235.1  OM670236.1  KT965208.1  KT965220.1 |
| *Grammostola quirogai* | KT965198.1  KT965199.1  KT965200.1  KT965201.1  KT965203.1  KT965204.1  KT965212.1  KT965213.1  KT965214.1  KT965215.1  KT965216.1  KT965217.1  KT965239.1  KT965251.1  KT965252.1  KT965253.1  KT965254.1  KT965264.1  KT965265.1  KT965268.1  KT965269.1  KT965273.1  KT965274.1  KT965275.1  KT965256.1  KT965270.1  KT965271.1  KT965272.1  KT965276.1  KT965234.1  KT965233.1  KT965229.1  KT965230.1  KT965231.1  KT965205.1  KT965221.1  KT965235.1  KT965240.1  KT965241.1  KT965242.1  KT965243.1  KT965244.1  KT965266.1 |
| *Grammostola rosea* | MK234708.1  MK234716.1  MK270580.1  MK270581.1  MK270583.1  KT022079.1  KT022081.1  KT022082.1  KT965262.1  KT965257.1  KT965260.1  KT965261.1  KT965259.1  MG273515.1 |
| *Grammostola vachoni* | OL906199.1  OL906198.1  OL906219.1  OL906215.1  OL906216.1  OL906217.1  OL906213.1  OL906212.1  OL906284.1  OL906203.1  OL906263.1  OL906280.1  OL906227.1  OL906269.1  OL906270.1  OL906283.1  OL906224.1  OL906266.1  OL906220.1  OL906218.1  OL906268.1  OL906226.1  OL906229.1  OL906252.1  OL906264.1  OL906265.1  OL906282.1  OL906262.1  OL906287.1  OL906225.1  OL906253.1  OL906274.1  OL906276.1  OL906249.1  OL906267.1  OL906258.1  OL906286.1  OL906237.1  OL906214.1  OL906250.1  OL906206.1  OL906271.1  OL906254.1  OL906256.1  OL906200.1  OL906272.1  OL906231.1  OL906201.1  OL906238.1  OL906236.1  OL906230.1  OL906241.1  OL906244.1  OL906246.1  OL906251.1  OL906273.1  OL906257.1  OL906275.1  OL906207.1  OL906285.1  OL906210.1  OL906233.1  OL906209.1  OL906247.1  OL906242.1  OL906211.1  OL906245.1  OL906255.1  OL906223.1  OL906202.1  OL906204.1  OL906208.1  OL906279.1  OL906234.1  OL906243.1  OL906232.1  OL906259.1  OL906239.1  OL906205.1  OL906240.1  OL906235.1  OL906278.1  OL906222.1  OL906248.1  OL906277.1  OL906260.1  OL906228.1  OL906221.1  OL906261.1 |
| *Haploclastus kayi* | MW462090.1  PV628768  PV628769 PV628770 PV628771 |
| *Ischnocolus elongatus* | OK428718.1 OK428719.1 OK428720.1 OK428721.1 OK428723.1 OK428724.1 OK428725.1 OK428726.1 OK428728.1 OK428729.1 OK428730.1 OK428731.1 OK428733.1 OK428734.1 OK428735.1 OK428736.1 OK428737.1 OK428738.1 OK428739.1 OK428740.1 OK428741.1 OK428742.1 OK428744.1 OK428747.1 OK428748.1 OK428751.1 OK428752.1 OK428753.1 OK428754.1 |
| *Ischnocolus mogadorensis* | OK428700  OK428703  OK428705  OK428711  OK428716  OK428722  OK428723  OK428724  OK428725  OK428727  OK428731  OK428732  OK428733  OK428734  OK428735  OK428737 |
| *Ischnocolus valentinus* | OK428729  OK428750  OK428751  OK428752  OK428753  OK428763  OK428766  OK428773  OK428774  OK428783  OK428784  OK428785  OK428786  OK428787  OK428788  OK428789  OK428790  OK428791 |
| *Lasiodora parahybana* | OK428718.1 MK234710.1 MK234713.1 MK234714.1 MK234722.1 MK234724.1 MK234725.1 MK270573.1 MK270574.1 |
| *Orphanaecus dichromatus* | PP726660.1 PP726661.1 PP726662.1 PP726663.1 PP726664.1 |
| *Plesiopelma longisternale* | KX835173.1 PP028765.1 PP726660.1 PP726661.1 PP726662.1 PP726663.1 PP726664.1 PP726659.1 PP726658.1 PP726657.1 PP726656.1 PP726655.1 PP726665.1 PP726666.1 |
| *Sericopelma melanosternum* | KX758187.1 KX758188.1 KX758189.1 KX758190.1 KR028321.1 |
| *Thrigmopoeus truculentus* | PV630228 PV630229 PV630230 PV630231 PV630232 PV630233 PV630234 PV630235 PV630236 PV630237 PV630238 PV630239 PV630240 PV630241 PV630242 PV630243 PV630244 PV630245 PV630246 PV630247 PV630248 PV630249 PV630250 PV630251 PV630252 PV630253 PV630254 PV630255 PV630256 PV630257 PV630258 |
| *Tiltocatl albopilosus* | MK213134.1 MK234707.1 MK234711.1 MK234719.1 MH362804.1 DQ224243.1 DQ224244.1 |
