## Supplementary material 3 for "Latitude Matters: A Global Phylogeographic Perspective on Climate-Driven Demographic Responses in Tarantulas"

**Figure S1:** Visualization of the predicted effects of the habitat suitability change on Tajima’ D for different latitudes. Plot (A) depicts results for the 10 percentile training presence and (B) equal training sensitivity and specificity. The predicted values along with the 95% confidence interval for every latitude class are plotted along with the observed values to compare model fit. This plot is for the 25-75% latitude binning scheme.

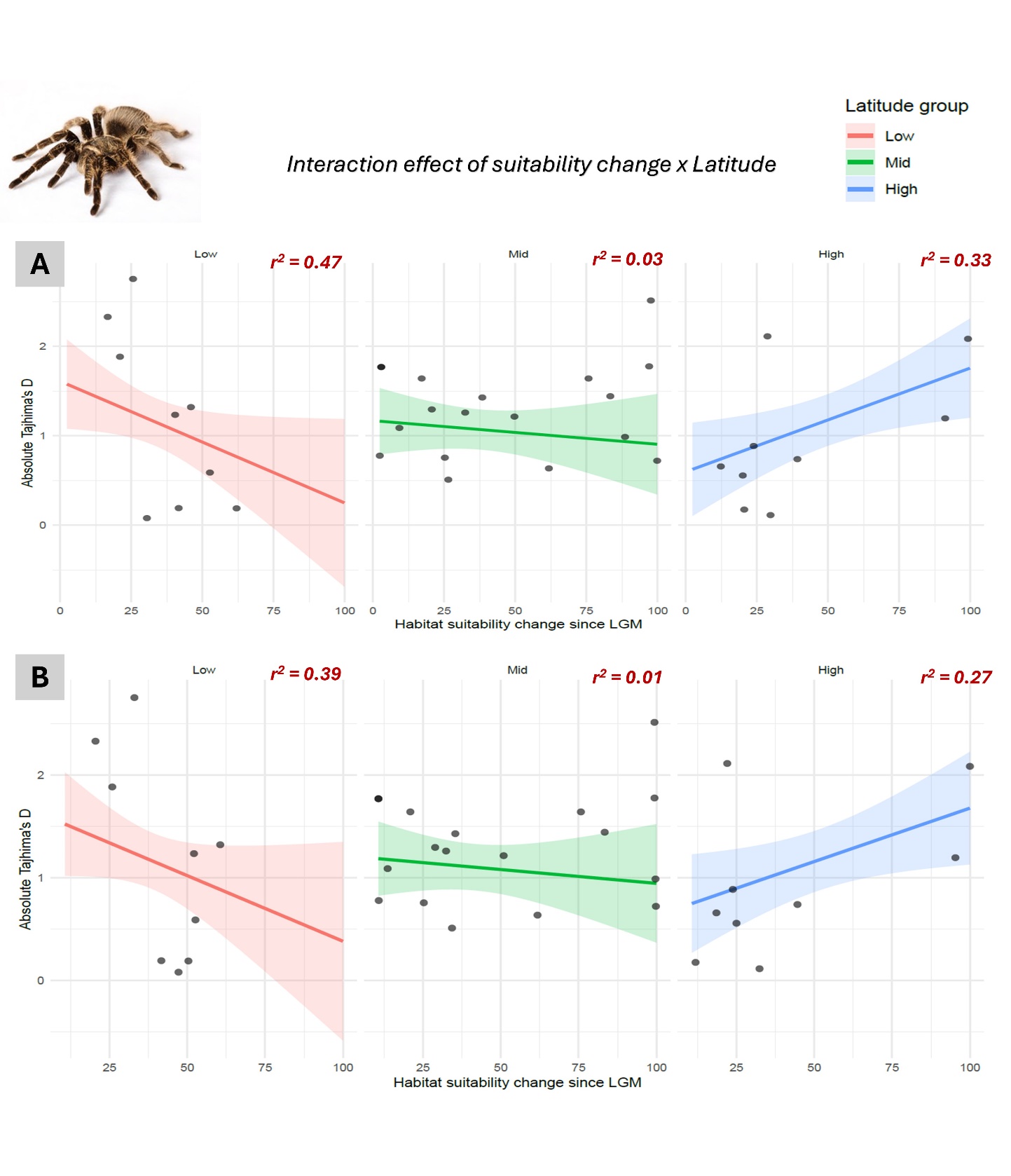

**Figure S2:** Visualization of the predicted effects of the habitat suitability change on Tajima’ D for different latitudes. Plot (A) depicts results for the 10 percentile training presence and (B) equal training sensitivity and specificity. The predicted values along with the 95% confidence interval for every latitude class are plotted along with the observed values to compare model fit. This plot is for the 35-65% latitude binning scheme.

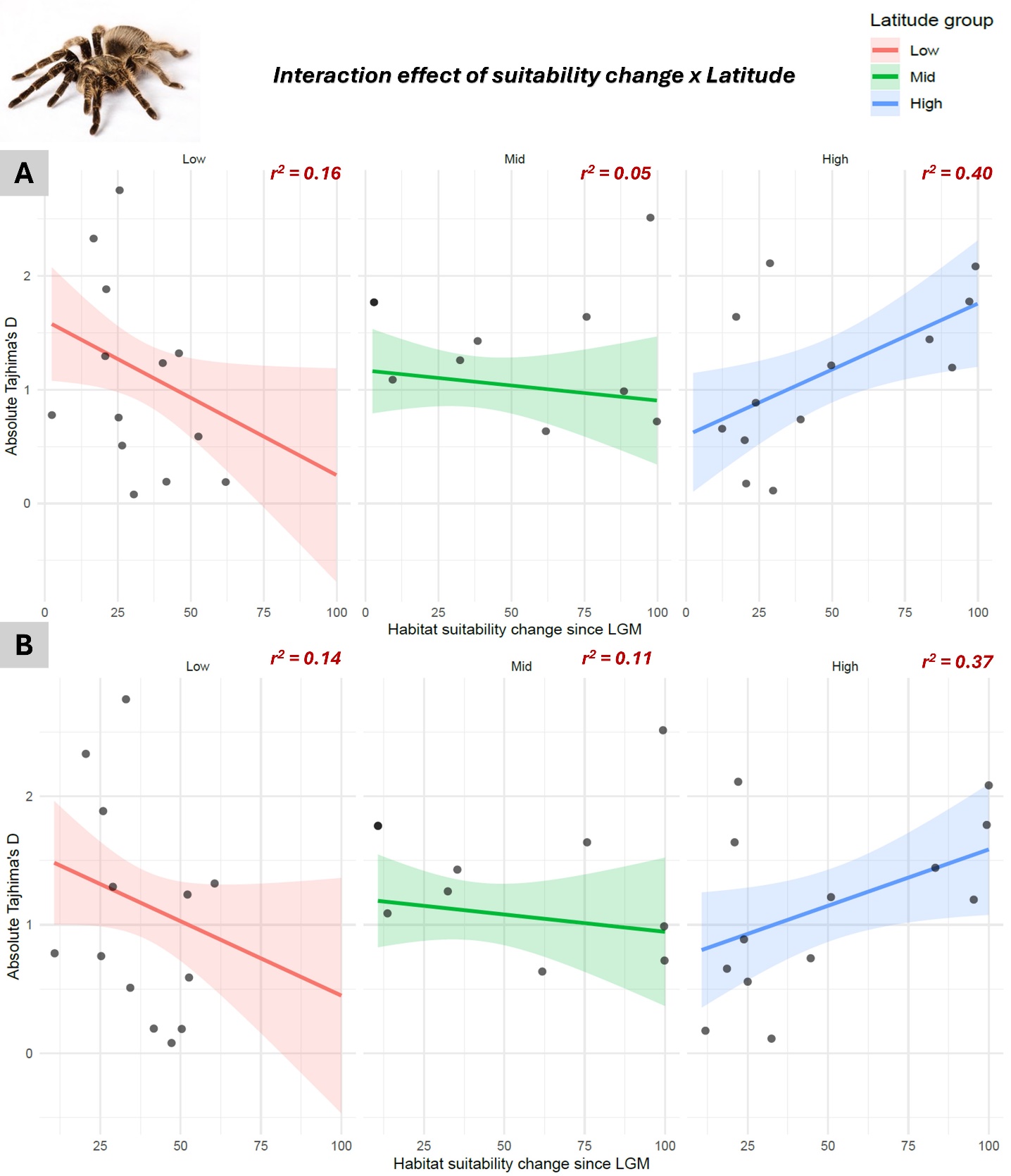
