## Supplementary material 4 for "Latitude Matters: A Global Phylogeographic Perspective on Climate-Driven Demographic Responses in Tarantulas"

| **Taxa** | **Present** | **LGM** |
| --- | --- | --- |
| *Aphonopelma anax* | 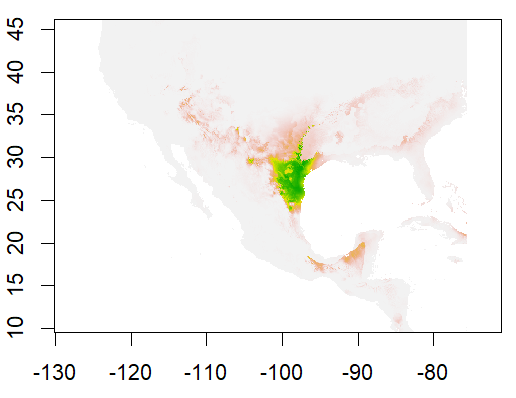 | 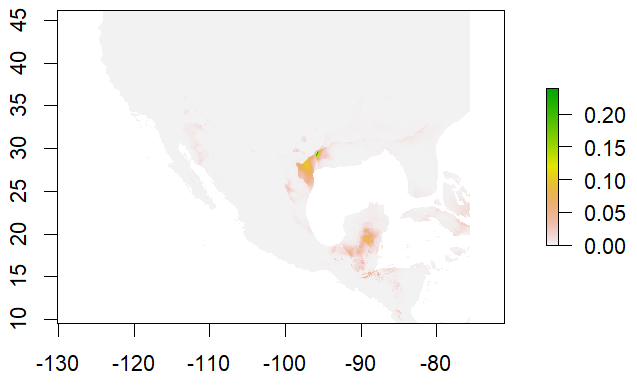 |
| *Aphonopelma armada* | 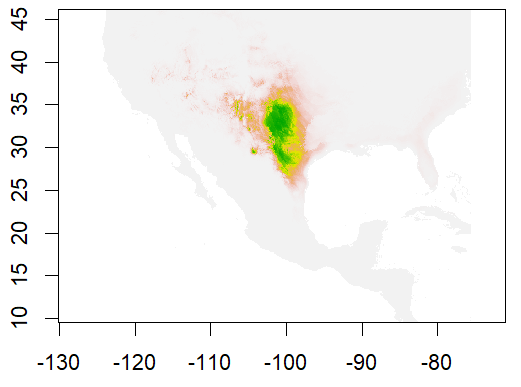 | 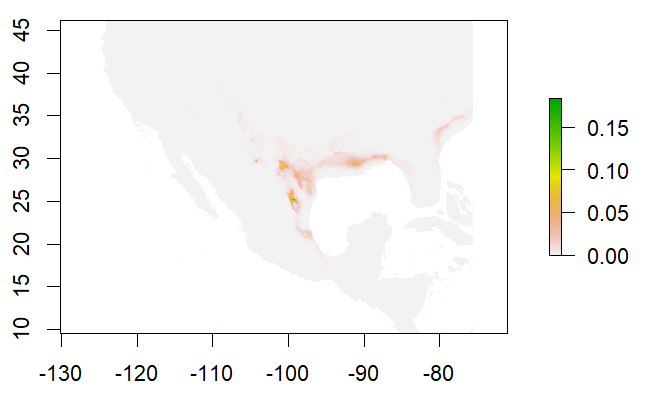 |
| *Aphonopelma chalcodes* | 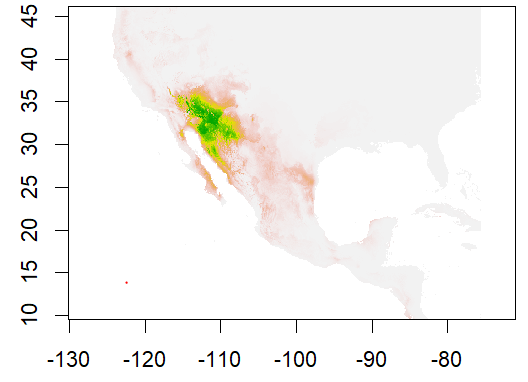 | 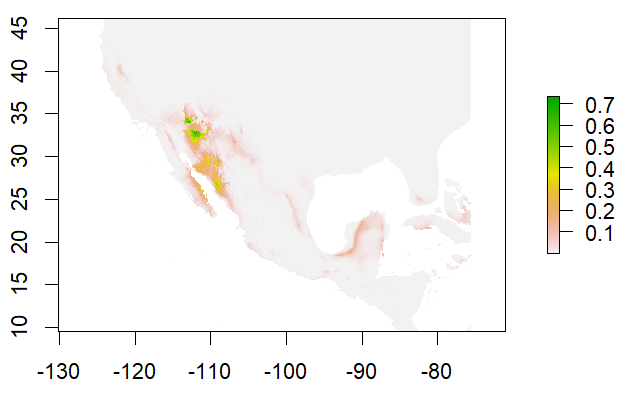 |
| *Aphonopelma eutylenum* | 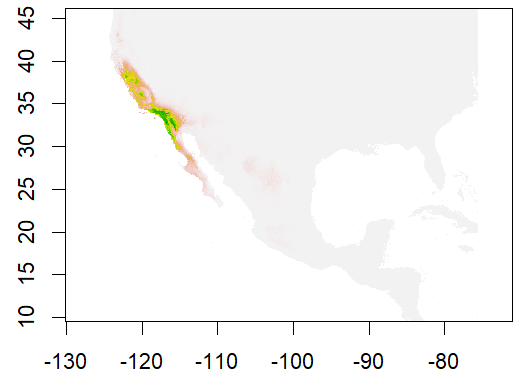 | 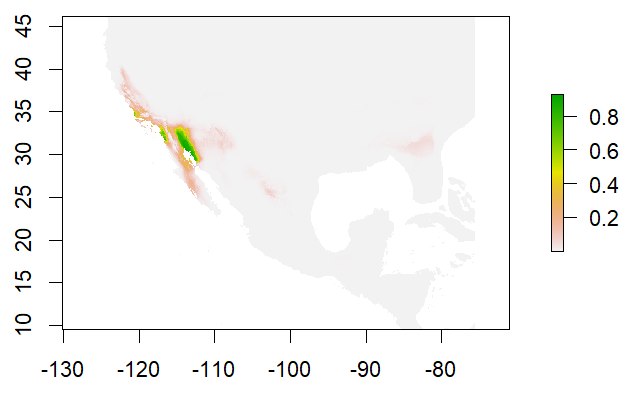 |
| *Aphonopelma gabeli* | 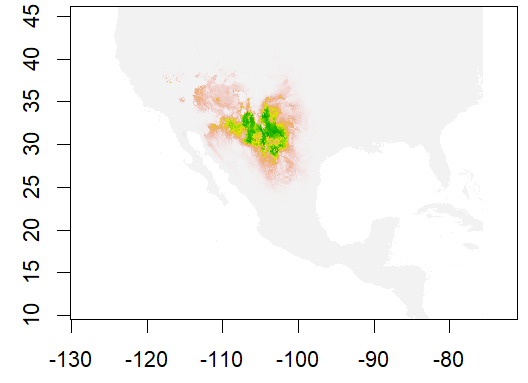 | 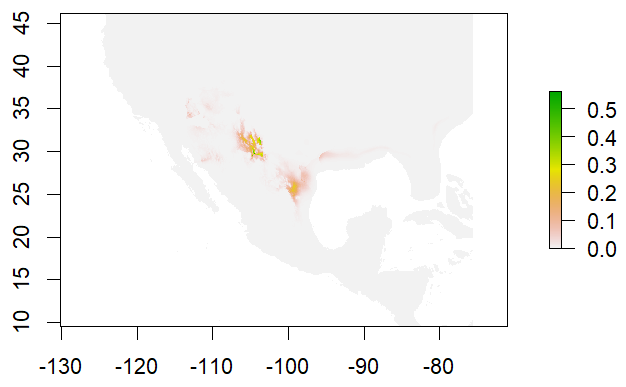 |
| *Aphonopelma hentzi* | 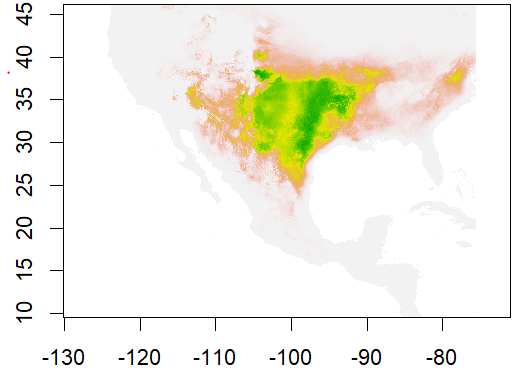 | 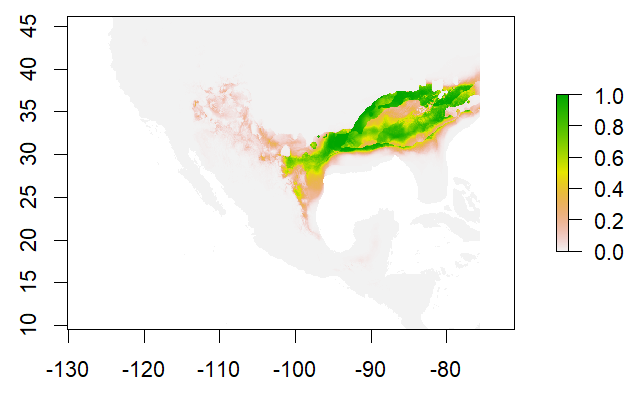 |
| *Aphonopelma iodius* | 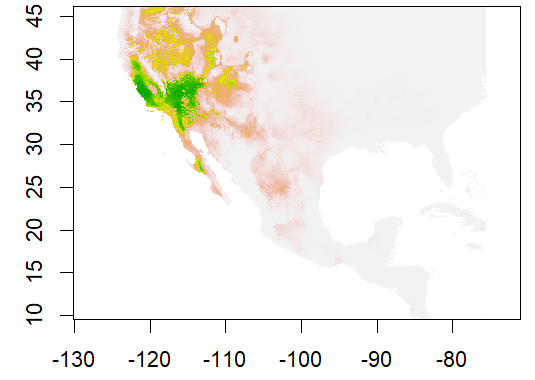 | 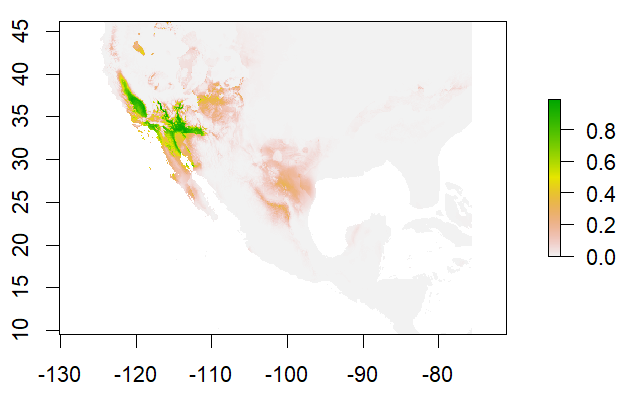 |
| *Aphonopelma joshua* | 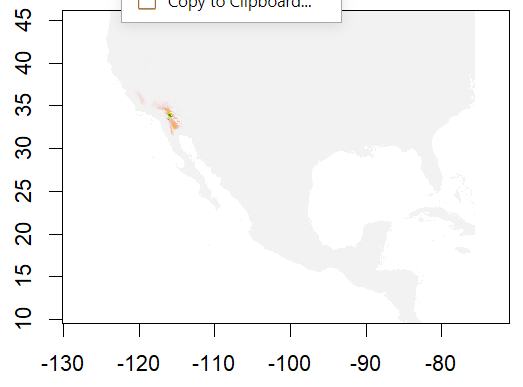 | 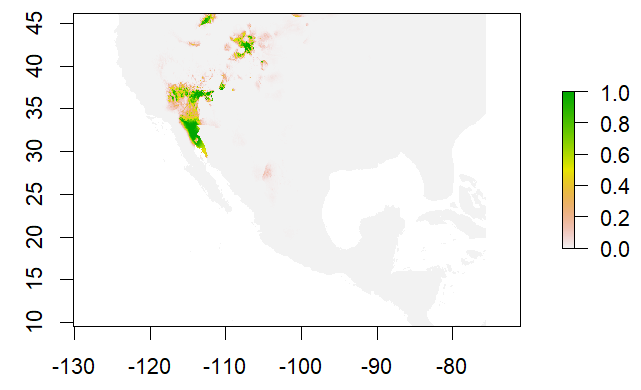 |
| *Aphonopelma marxi* | 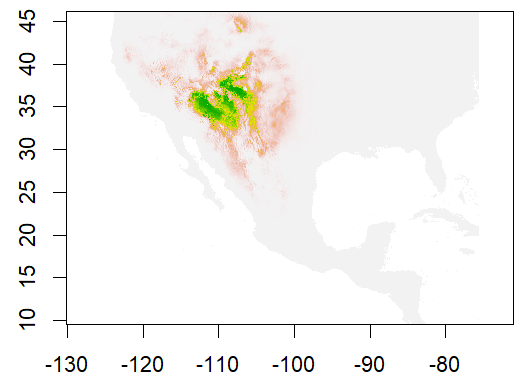 | 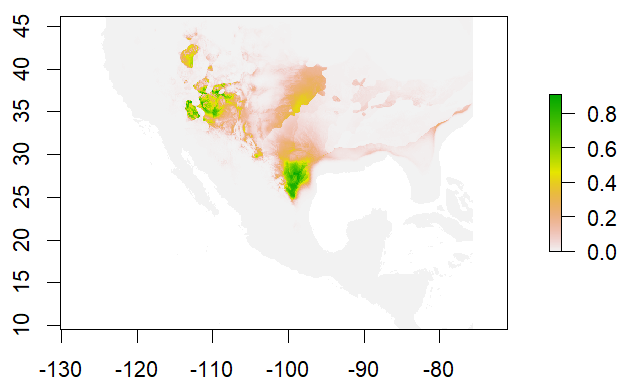 |
| *Aphonopelma moderatum* | 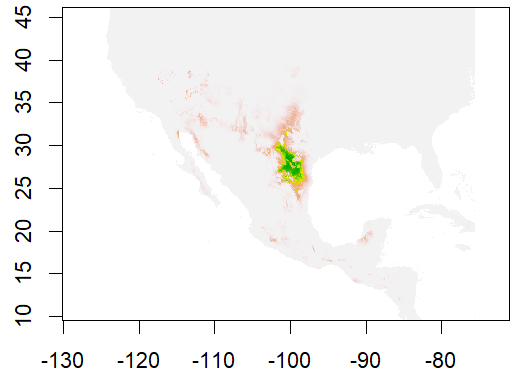 | 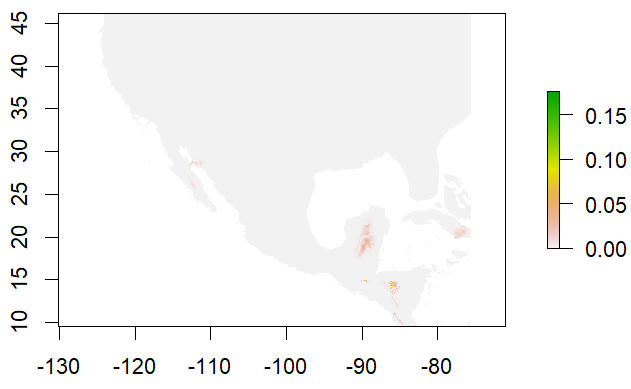 |
| *Aphonopelma paloma* | 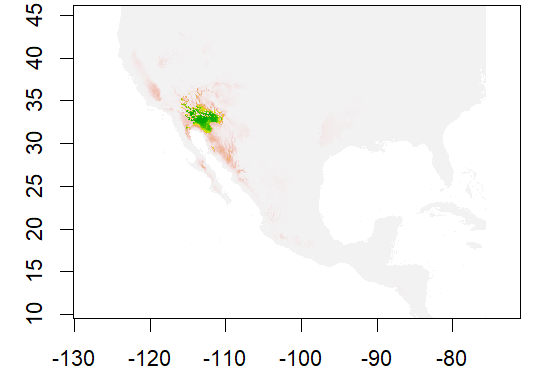 | 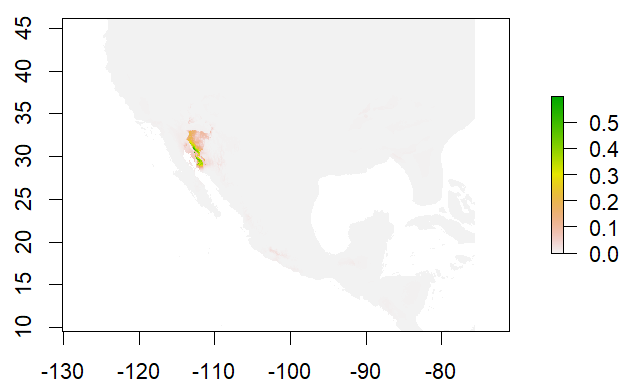 |
| *Aphonopelma steindechneri* | 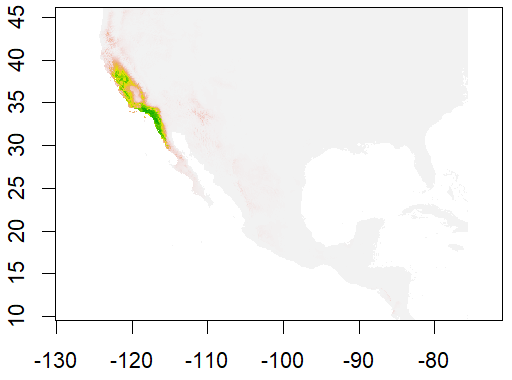 | 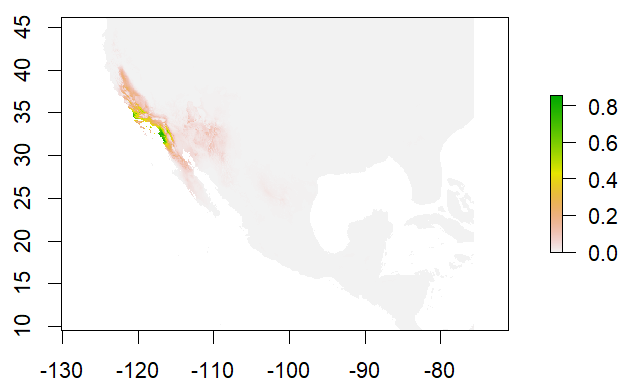 |
| *Aphonopelma vorhiesi* | 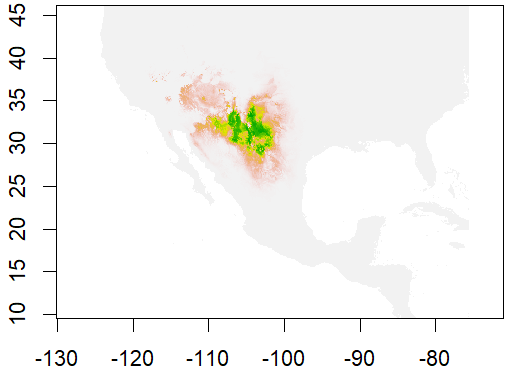 | 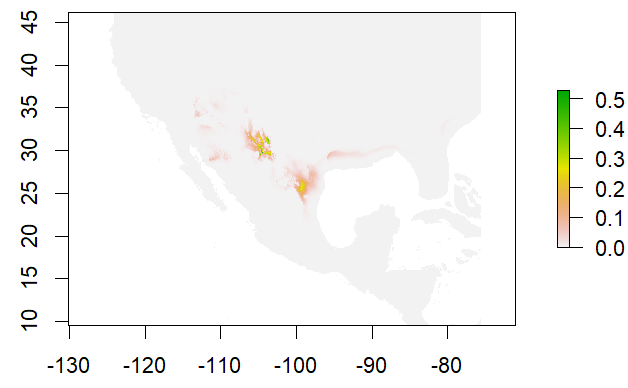 |
| *Bonnetina avaie* | 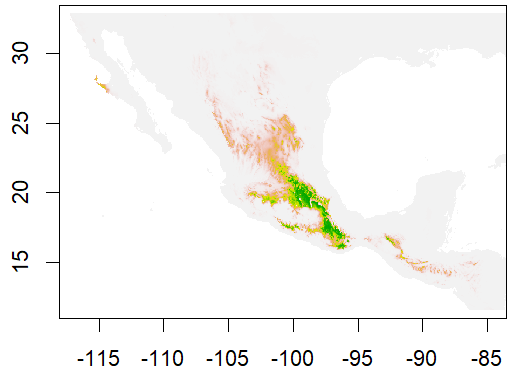 | 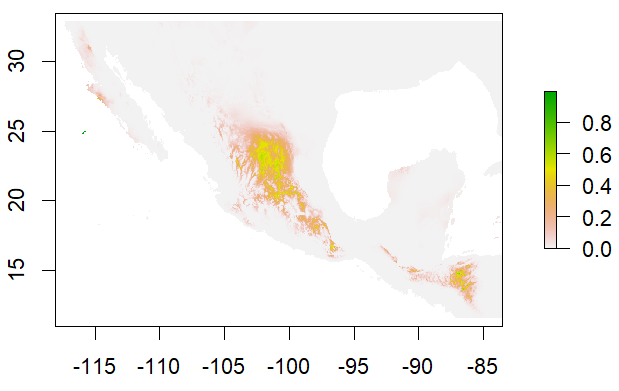 |
| *Bonnetina cyaneifemur* | 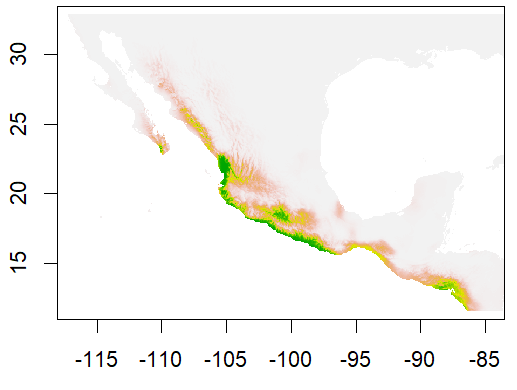 | 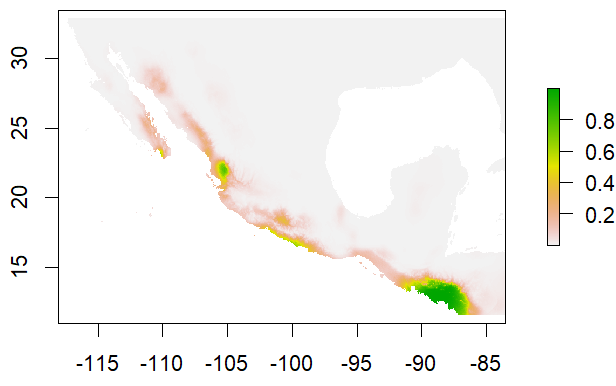 |
| *Bonnetina papalutlensis* | 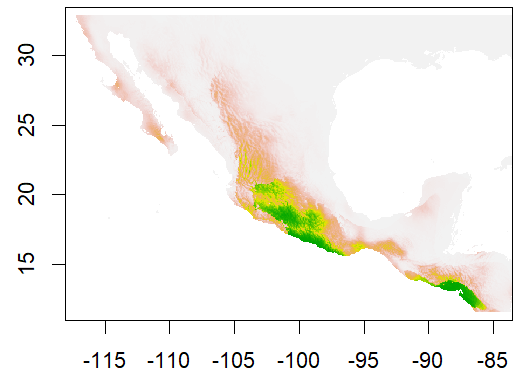 | 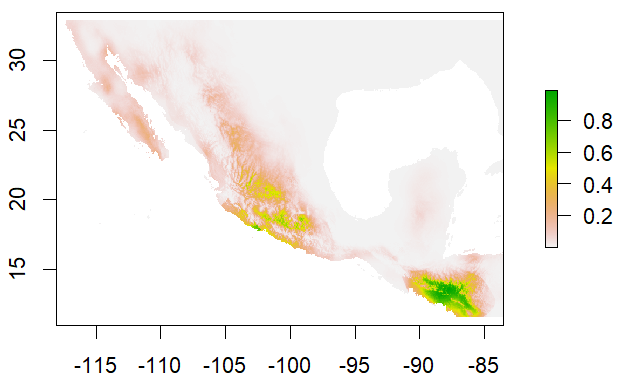 |
| *Brachypelma klassi* | 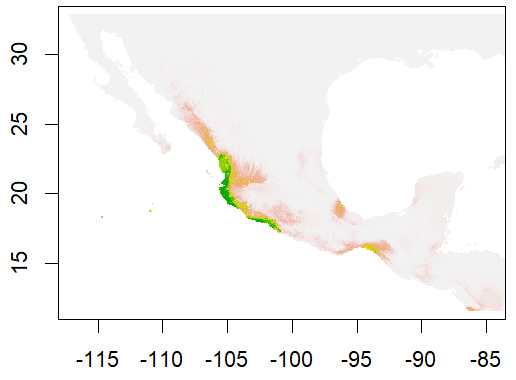 | 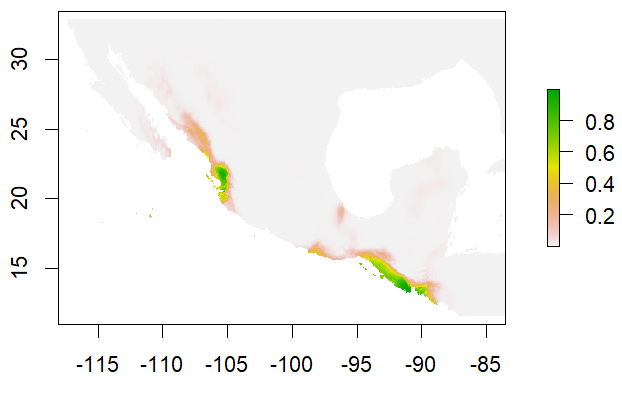 |
| *Brachypelma albiceps* | 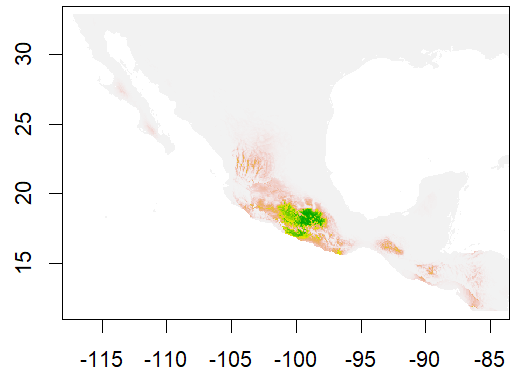 | 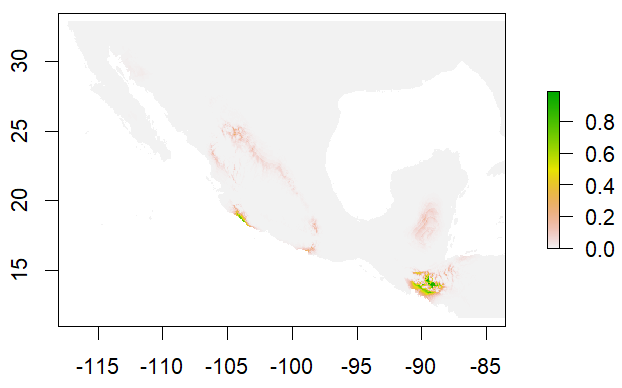 |
| *Brachypelma baumgarteni* | 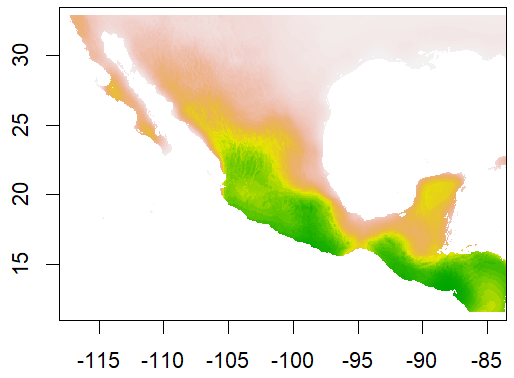 | 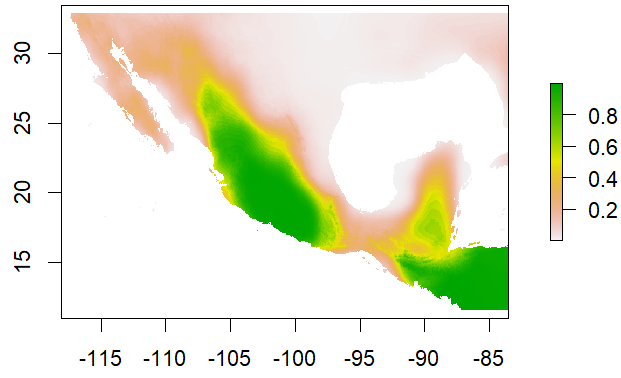 |
| *Brachypelma boehmei* | 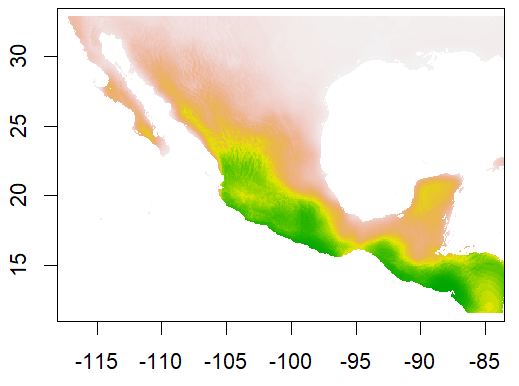 | 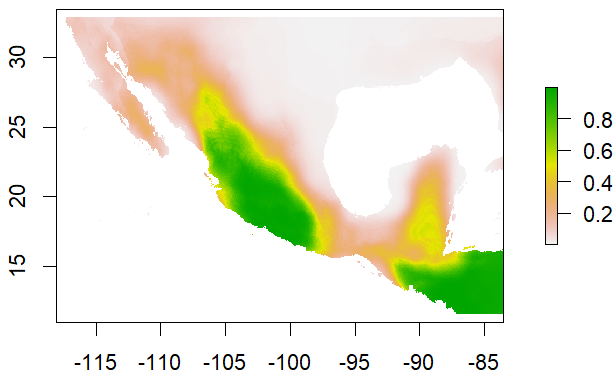 |
| *Brachypelma hamorii* | 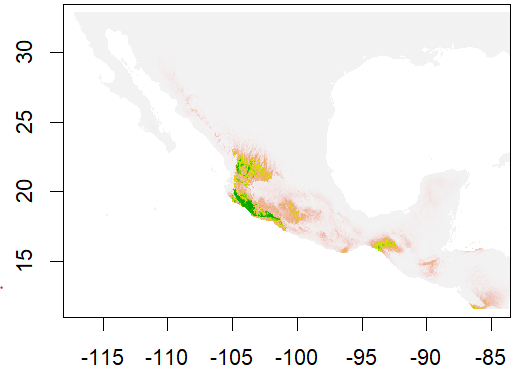 | 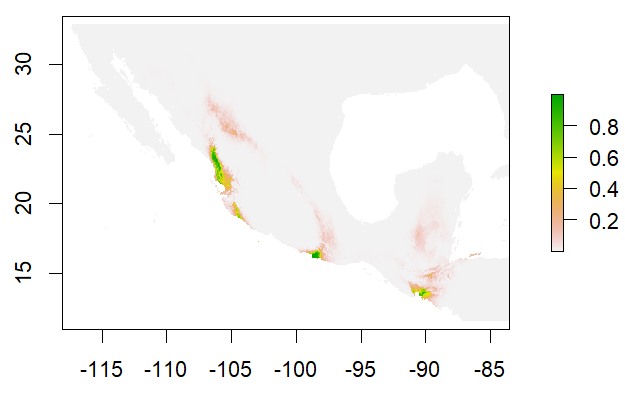 |
| *Brachypelma kahlenbergii* | 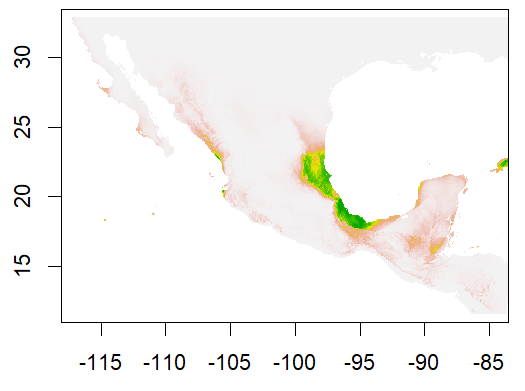 | 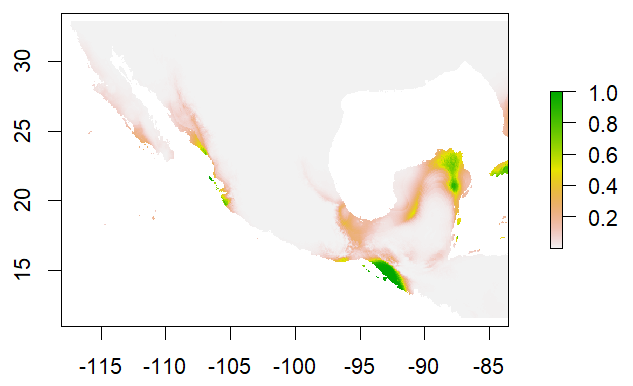 |
| *Brachypelma klassi* | 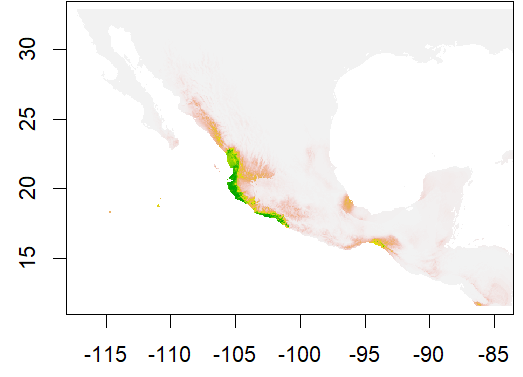 | 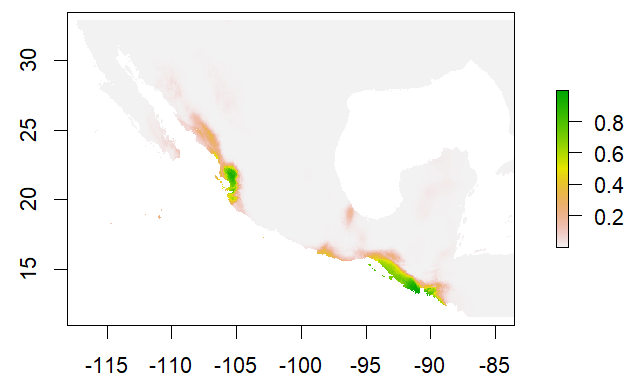 |
| *Brachypelma schroederi* | 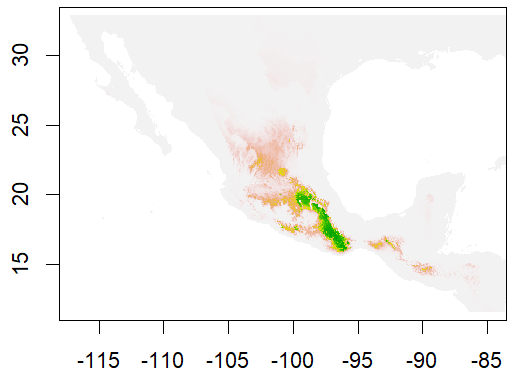 | 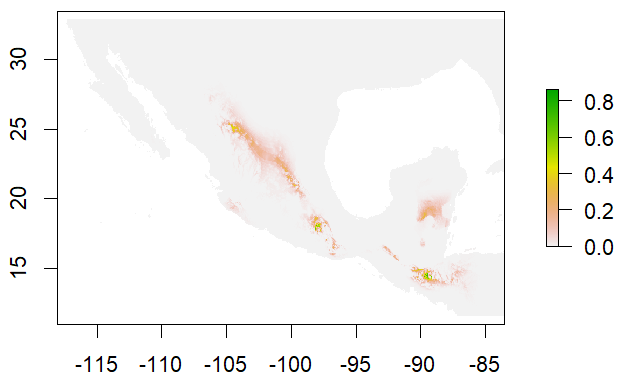 |
| *Brachypelma smithi* | 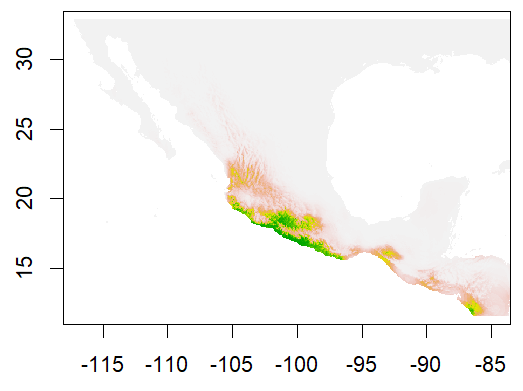 | 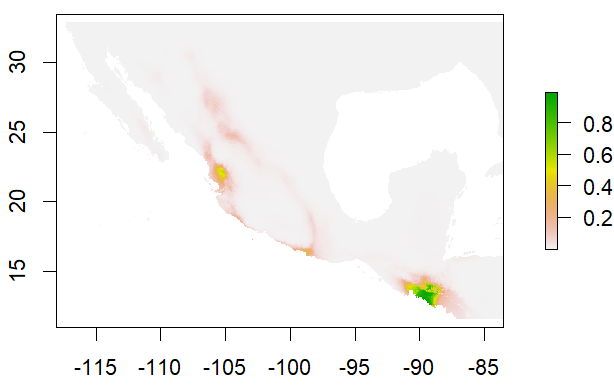 |
| *Brachypelma vagans* | 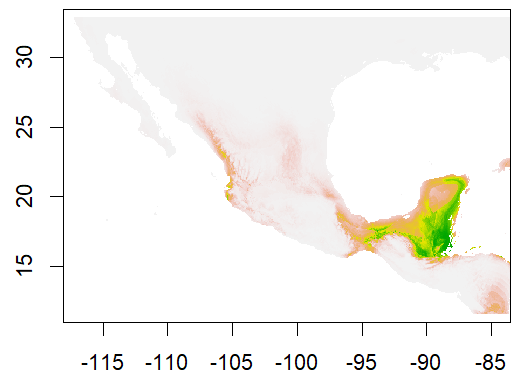 | 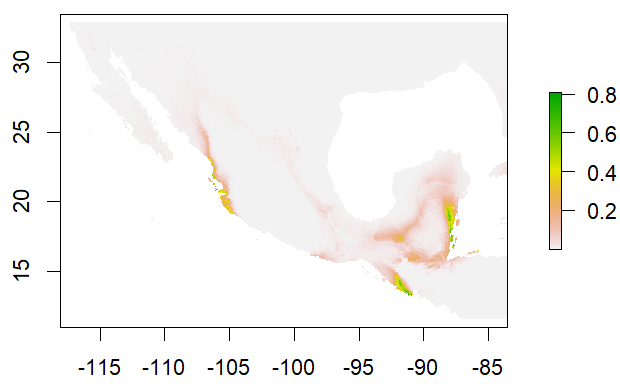 |
| *Brachypelma verdezi* | 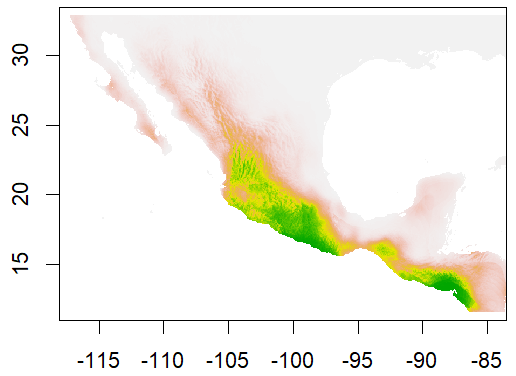 | 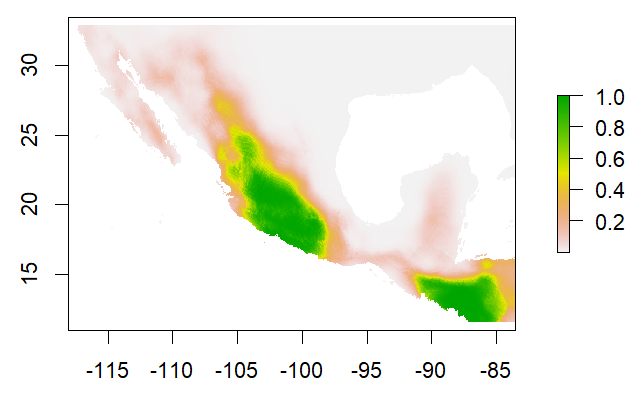 |
| *Grammostola anthracina* | 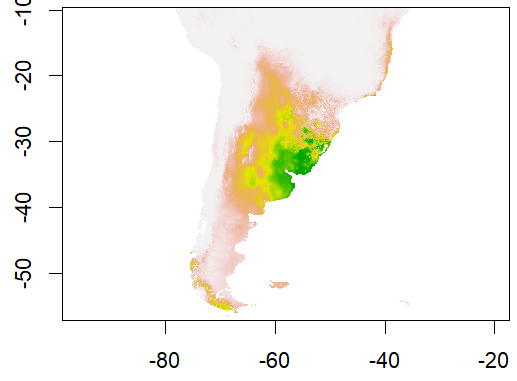 | 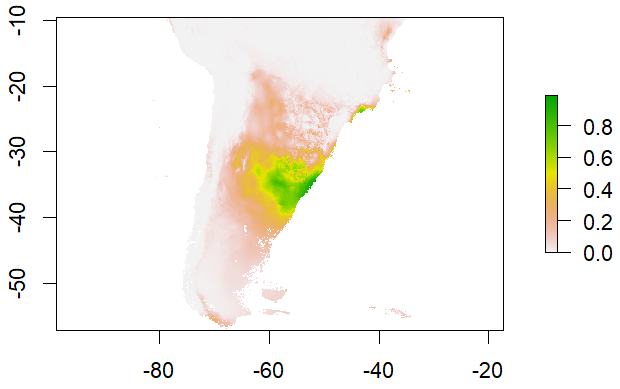 |
| *Grammostola quirogai* | 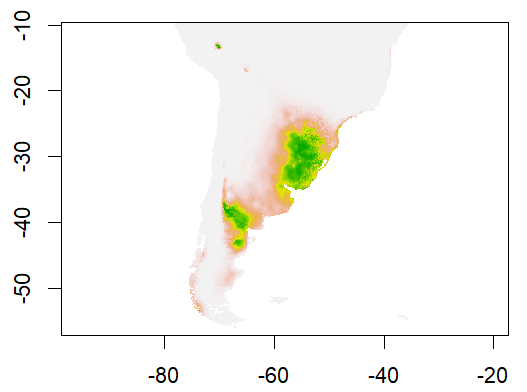 | 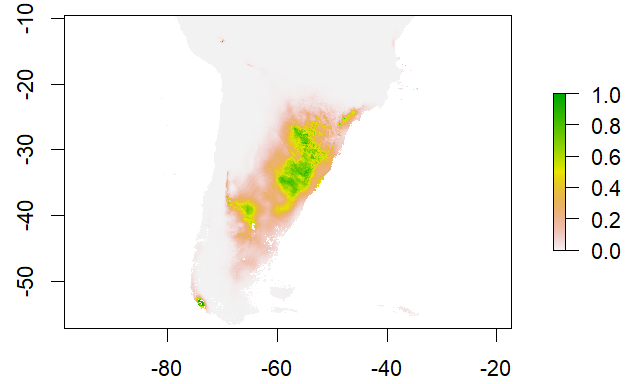 |
| *Grammostola rosea* | 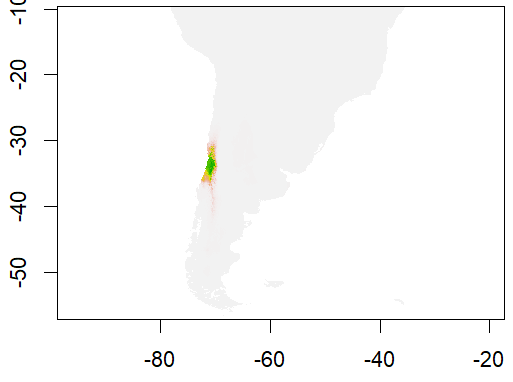 | 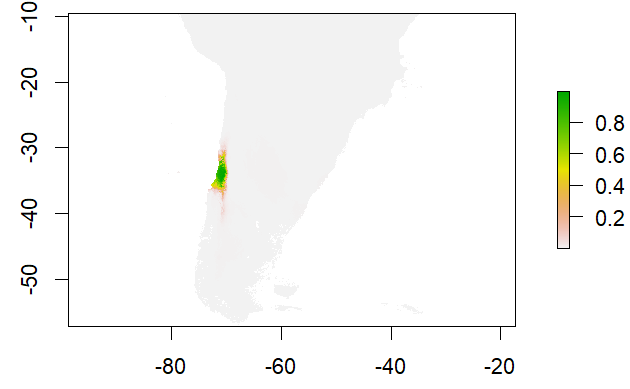 |
| *Grammostola vachoni* | 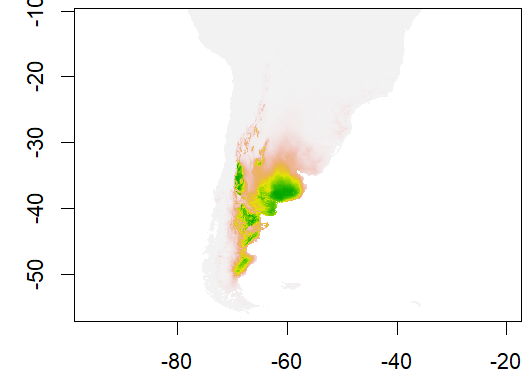 | 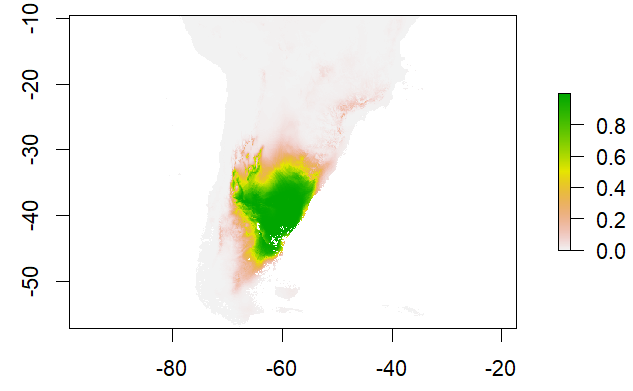 |
| *Haploclastus kayi* | 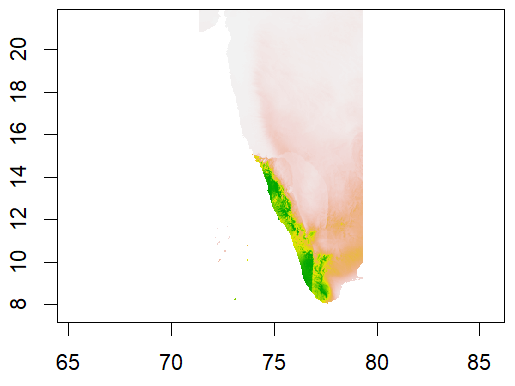 | 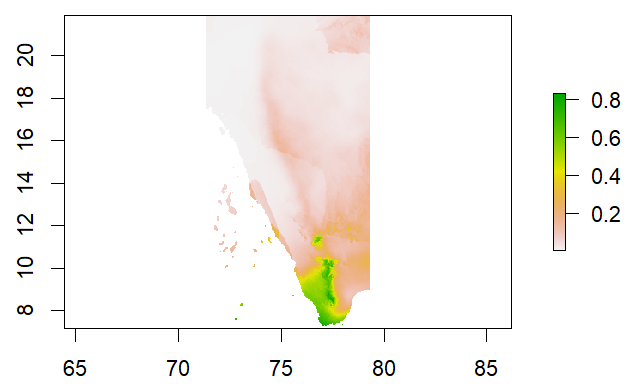 |
| *Ischnocolus elongatus* | 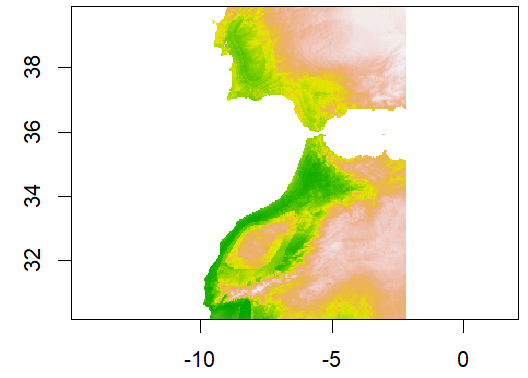 | 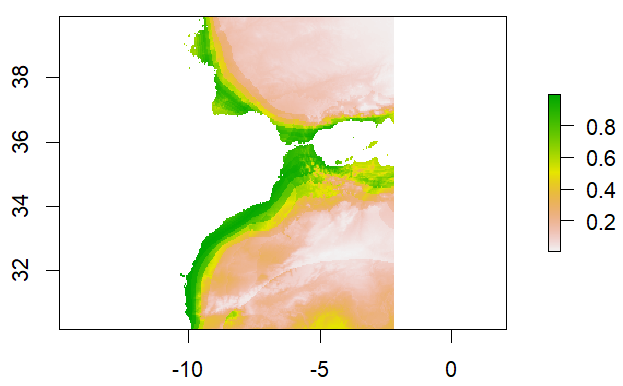 |
| *Lasiodora parahybana* | 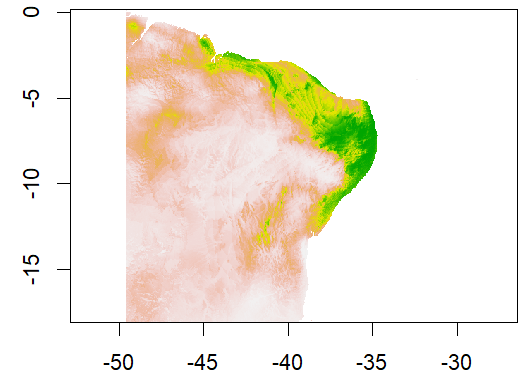 | 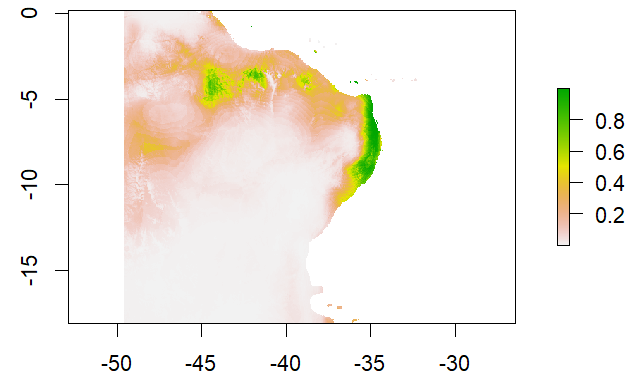 |
| *Plesiopelma longisternale* | 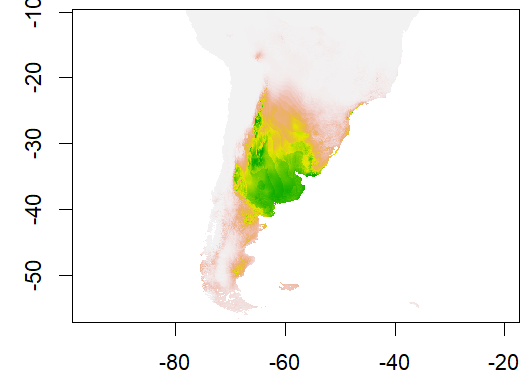 | 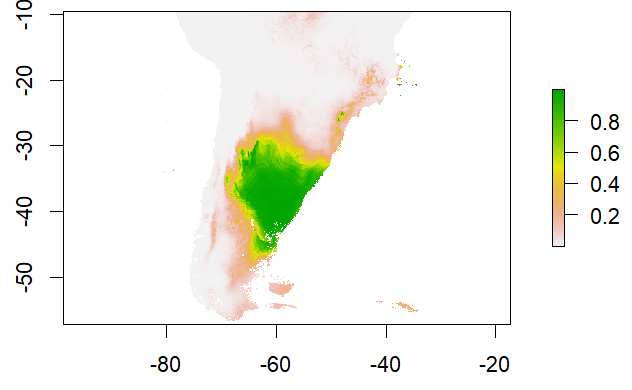 |
| *Thrigmopoeus truculentus* | 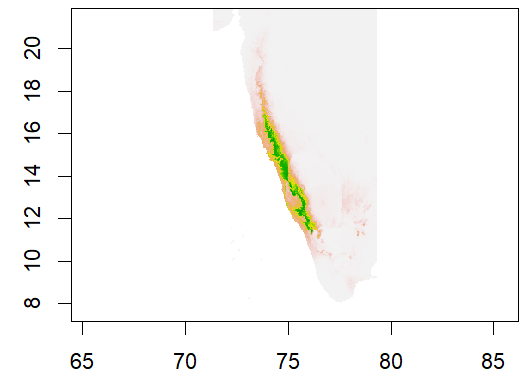 | 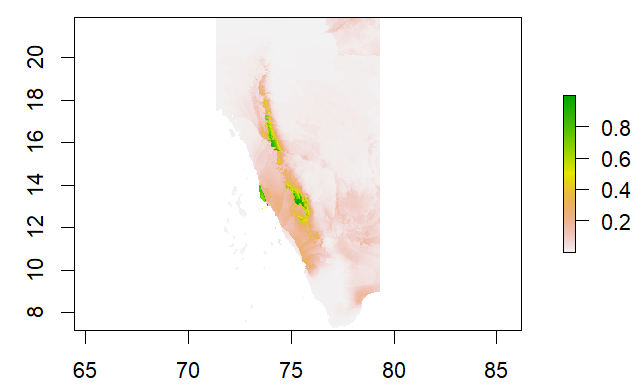 |
| *Tiltocatl albopilosum* | 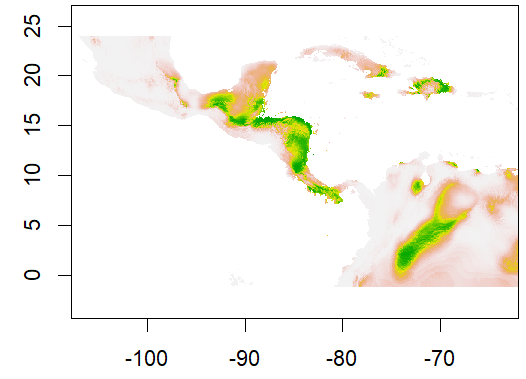 | 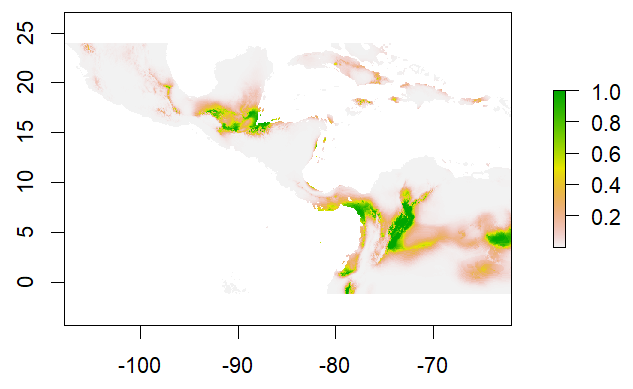 |
