## Supplementary material 5 for "Latitude Matters: A Global Phylogeographic Perspective on Climate-Driven Demographic Responses in Tarantulas": glm_script.R

```
library(MuMIn)

#importing the data and removing na values#####

data<-read.csv("data glm.csv")
data

data<-na.omit(data)
data


#######################################################################################################################################################################################
########GLM with demographic response (Tajhima's D) as response variable and latitude and suitability change (minimum training presence cut off) as predictor variable################
#####################################################################################################################################################################################


model1<-glm(abs_Tajhima.s_D~ abs.lat, data = data, family = gaussian)
summary(model1)

model2<-glm(abs_Tajhima.s_D~ Percent_affected_area_a, data = data, family = gaussian)
summary(model2)

model3<-glm(abs_Tajhima.s_D~ Percent_affected_area_a * abs.lat, data = data, family = gaussian)
summary(model3)

AIC(model1, model2, model3)
model_selection <- model.sel(model1, model2, model3, rank = "AICc")
print(model_selection)
write.csv(model_selection,"table 1.csv")

###plotting the model####

library(ggplot2)

# Step 1: Create a prediction grid
new_data <- expand.grid(
  Percent_affected_area_a = seq(min(data$Percent_affected_area_a, na.rm = TRUE),
                                       max(data$Percent_affected_area_a, na.rm = TRUE),
                                       length.out = 100),
  abs.lat = quantile(data$abs.lat, probs = c(0.25, 0.5, 0.75), na.rm = TRUE)  # low, mid, high
)

# Step 2: Predict with standard errors for CI bands
pred <- predict(model3, newdata = new_data, se.fit = TRUE)

# Add predicted values and 95% confidence intervals to new_data
new_data$fit <- pred$fit
new_data$se <- pred$se.fit
new_data$lower <- new_data$fit - 1.96 * new_data$se
new_data$upper <- new_data$fit + 1.96 * new_data$se


###############3 separate plots####################

# First, get the actual quantile values
lat_quantiles <- quantile(data$abs.lat, probs = c(0.35, 0.5, 0.65), na.rm = TRUE)
lat_quantiles

# Add lat_bin to your data
new_data$lat_bin <- cut(
  new_data$abs.lat,
  breaks = c(-Inf, lat_quantiles[1], lat_quantiles[3], Inf),
  labels = c("Low", "Mid", "High"),
  right = TRUE,
  include.lowest = TRUE
)


ggplot() +
  geom_ribbon(data = new_data, aes(x = Percent_affected_area_a, ymin = lower, ymax = upper, fill = lat_bin), alpha = 0.2) +
  geom_line(data = new_data, aes(x = Percent_affected_area_a, y = fit, color = lat_bin), size = 1.2) +
  geom_point(data = data, aes(x = Percent_affected_area_a, y = abs_Tajhima.s_D), size = 2, alpha = 0.6, color = "black") +
  facet_wrap(~lat_bin) +
  labs(
    x = "Habitat suitability change since LGM",
    y = "Absolute Tajhima's D",
    title = "Interaction effect of Habitat Change � Latitude",
    color = "Latitude group",
    fill = "Latitude group"
  ) +
  theme_minimal()


#######################################################################################################################################################################################
########GLM with demographic response (Tajhima's D) as response variable and latitude and suitability change (equal sensitivity and specificity cut off) as predictor variable################
#####################################################################################################################################################################################

model1<-glm(abs_Tajhima.s_D~ abs.lat, data = data, family = gaussian)
summary(model1)

model2<-glm(abs_Tajhima.s_D~ Percent_affected_area_b, data = data, family = gaussian)
summary(model2)

model3<-glm(abs_Tajhima.s_D~ Percent_affected_area_b * abs.lat, data = data, family = gaussian)
summary(model3)

AIC(model1, model2, model3)
model_selection <- model.sel(model1, model2, model3, rank = "AICc")
print(model_selection)
write.csv(model_selection,"table 2.csv")

###plotting the model####

library(ggplot2)

# Step 1: Create a prediction grid
new_data <- expand.grid(
  Percent_affected_area_b = seq(min(data$Percent_affected_area_b, na.rm = TRUE),
                                max(data$Percent_affected_area_b, na.rm = TRUE),
                                length.out = 100),
  abs.lat = quantile(data$abs.lat, probs = c(0.35, 0.5, 0.65), na.rm = TRUE)  # low, mid, high
)

# Step 2: Predict with standard errors for CI bands
pred <- predict(model3, newdata = new_data, se.fit = TRUE)

# Add predicted values and 95% confidence intervals to new_data
new_data$fit <- pred$fit
new_data$se <- pred$se.fit
new_data$lower <- new_data$fit - 1.96 * new_data$se
new_data$upper <- new_data$fit + 1.96 * new_data$se


###############3 separate plots####################

# First, get the actual quantile values
lat_quantiles <- quantile(data$abs.lat, probs = c(0.35, 0.5, 0.65), na.rm = TRUE)

# Add lat_bin to your original data
data$lat_bin <- cut(
  data$abs.lat,
  breaks = c(-Inf, lat_quantiles[1], lat_quantiles[3], Inf),
  labels = c("Low", "Mid", "High"),
  right = TRUE,
  include.lowest = TRUE
)

new_data$lat_bin <- factor(new_data$abs.lat,
                           levels = lat_quantiles,
                           labels = c("Low", "Mid", "High"))


ggplot() +
  geom_ribbon(data = new_data, aes(x = Percent_affected_area_b, ymin = lower, ymax = upper, fill = lat_bin), alpha = 0.2) +
  geom_line(data = new_data, aes(x = Percent_affected_area_b, y = fit, color = lat_bin), size = 1.2) +
  geom_point(data = data, aes(x = Percent_affected_area_b, y = abs_Tajhima.s_D), size = 2, alpha = 0.6, color = "black") +
  facet_wrap(~lat_bin) +
  labs(
    x = "Habitat suitability change since LGM",
    y = "Absolute Tajhima's D",
    title = "Interaction effect of Habitat Change � Latitude",
    color = "Latitude group",
    fill = "Latitude group"
  ) +
  theme_minimal()


#######################################################################################################################################################################################
########GLM with nucleotide diversity as response variable and latitude and suitability change (equal sensitivity and specificity cut off) as predictor variable################
#####################################################################################################################################################################################

model1<-glm(Nucleotide_diversity~ abs.lat, data = data, family = gaussian)
summary(model1)

model2<-glm(Nucleotide_diversity~ Percent_affected_area_b, data = data, family = gaussian)
summary(model2)

model3<-glm(Nucleotide_diversity~ Percent_affected_area_b * abs.lat, data = data, family = gaussian)
summary(model3)

model_selection <- model.sel(model1, model2, model3, rank = "AICc")
print(model_selection)
write.csv(model_selection,"table 3.csv")


#######################################################################################################################################################################################
########GLM with nucleotide diversity as response variable and latitude and suitability change (minimum training presence) as predictor variable################
#####################################################################################################################################################################################

model1<-glm(Nucleotide_diversity~ abs.lat, data = data, family = gaussian)
summary(model1)

model2<-glm(Nucleotide_diversity~ Percent_affected_area_a, data = data, family = gaussian)
summary(model2)

model3<-glm(Nucleotide_diversity~ Percent_affected_area_a * abs.lat, data = data, family = gaussian)
summary(model3)

AIC(model1, model2, model3)
model_selection <- model.sel(model1, model2, model3, rank = "AICc")
print(model_selection)
write.csv(model_selection,"table 4.csv")


#######################################################################################################################################################################################
########GLM with suitability change as response variable and abs latitude as predictor variable################
#####################################################################################################################################################################################

model1<-glm(Percent_affected_area_a~ abs.lat, data = data, family = gaussian)
summary(model1)

model2<-glm(Percent_affected_area_b~ abs.lat, data = data, family = gaussian)
summary(model2)


#######################################################################################################################################################################################
########Plot Tajhima's D values################
#####################################################################################################################################################################################
library(ggplot2)
library(dplyr)

data <- data %>%
  filter(!is.na(Tajhima.s.D.value)) %>%
  mutate(Taxa = factor(Taxa, levels = Taxa[order(Tajhima.s.D.value)]),
         Tajima_sign = ifelse(Tajhima.s.D.value >= 0, "Positive", "Negative"))


# Plot
ggplot(data, aes(x = Taxa, y = Tajhima.s.D.value, fill = Tajima_sign)) +
  geom_bar(stat = "identity") +
  scale_fill_manual(values = c("Positive" = "tomato", "Negative" = "steelblue")) +
  coord_flip() +
  theme_minimal() +
  labs(title = "Tajima's D values by Species",
       x = "Taxa",
       y = "Tajhima.s.D.value",
       fill = "Tajima's D Sign") +
  theme(axis.text.y = element_text(size = 10),
        plot.title = element_text(hjust = 0.5, face = "bold"))


#######################################################################################################################################################################################
########Plot Nucleotide diversity################
#####################################################################################################################################################################################

# Clean and prepare the data
data <- data %>%
  filter(!is.na(Nucleotide_diversity), !is.na(Nature_of_change)) %>%
  mutate(Taxa = factor(Taxa, levels = Taxa[order(Nucleotide_diversity)]))

# Create the bar plot
ggplot(data, aes(x = Taxa, y = Nucleotide_diversity, fill = Nature_of_change)) +
  geom_bar(stat = "identity") +
  coord_flip() +
  theme_minimal() +
  labs(title = "Nucleotide Diversity by Species",
       x = "Species",
       y = "Nucleotide_diversity",
       fill = "Nature_of_change") +
  theme(axis.text.y = element_text(size = 10),
        plot.title = element_text(hjust = 0.5, face = "bold")) +
  scale_fill_brewer(palette = "Set2")  # Optional: prettier colors


#######################################################################################################################################################################################
########Plot percent area affected under different cut-offs for comparison################
#####################################################################################################################################################################################

library(ggplot2)
library(dplyr)
library(tidyr)

# Reorder species by Percent_affected_area_a
data <- data %>%
  arrange(desc(Percent_affected_area_a)) %>%
  mutate(Taxa = factor(Taxa, levels = unique(Taxa)))


# Reshape for plotting
long_data <- data %>%
  select(Taxa, Percent_affected_area_a, Percent_affected_area_b) %>%
  pivot_longer(cols = starts_with("Percent_affected_area"),
               names_to = "Area_Type",
               values_to = "Percent_Affected") %>%
  filter(!is.na(Percent_Affected))


ggplot(long_data, aes(x = Taxa, y = Percent_Affected, fill = Area_Type)) +
  geom_bar(stat = "identity", position = position_dodge(width = 0.8)) +
  coord_flip() +
  theme_minimal() +
  labs(title = "Comparison of Affected Areas by Species",
       x = "Species",
       y = "Percent Affected Area",
       fill = "Area Type") +
  theme(axis.text.y = element_text(size = 9),
        plot.title = element_text(hjust = 0.5, face = "bold")) +
  scale_fill_manual(values = c("Percent_affected_area_a" = "skyblue",
                               "Percent_affected_area_b" = "orange"))
```
