## Supplementary material 5 for "Latitude Matters: A Global Phylogeographic Perspective on Climate-Driven Demographic Responses in Tarantulas": spatial_data_processing_script.R

```
library(raster)
library(rgdal)
library(dismo)
library(maptools)
library(sp)

thrigmopoeus<-read.csv("all distribution data.csv")
thrigmopoeus

data(wrld_simpl)
plot(wrld_simpl, xlim=c(75,76), ylim=c(6,20), axes=TRUE, col="light yellow")

# restore the box around the map
box()

# add the points
points(thrigmopoeus$long, thrigmopoeus$lat, col='orange', pch=20, cex=0.75)


wodups <- duplicated(thrigmopoeus[, c('lat', 'long')])
wodups
# keep the records that are _not_ duplicated
final <- thrigmopoeus[!wodups, ]
final

final_data<-cbind(final$lat, final$long)
final_data

# create a RasterLayer with the extent of distribution
thrigmopoeus<-thrigmopoeus[,-1]

coordinates(thrigmopoeus) = ~long+lat # Convert dataframe to SpatialPoints object
thrigmopoeus
crs(thrigmopoeus) = crs(wrld_simpl) # Associate coordinate reference system
class(thrigmopoeus) # Check the class
ovr = over(thrigmopoeus, wrld_simpl) # help(over)
class(ovr); names(over) # a dataframe, check the column names
head(ovr)
country = ovr$NAME # Country names associated with locations
sum(is.na(country)) # Number of rows with countries = NA
is_ocean = which(is.na(country)); head(is_ocean); table(is_ocean) # Check the variable is_ocean
lonlat_ocean = thrigmopoeus@coords[is_ocean,] # lon-lat of points which fall into ocean
lonlat_ocean
thrigmopoeus@coords = thrigmopoeus@coords[-is_ocean,] # Remove those coordinates
plot(thrigmopoeus, col = "green", cex= 0.5, pch=16) # Plot all points with green
plot(wrld_simpl, add=TRUE, lwd = 2) # Add the world map, note they have same crs
points(lonlat_ocean, col="red", cex = 0.5, pch=16, lwd= 2) # Ocean points marked as red
box() # Bounding box


r<-raster(thrigmopoeus)
r
res(r) <- 0.1
r
r <- extend(r, extent(r)+1)
r
acsel <- gridSample(thrigmopoeus, r, n=1)
acsel
p <- rasterToPolygons(r)
plot(p, border='gray')
points(thrigmopoeus)
points(acsel, cex=0.5, col='red', pch='x')


write.csv(acsel,"cleaned data.csv")


####removing correlation from present day predictor bioclimatic variables###############
setwd("D:/PhD works/world clim rasters/clipped rasters India")


library(raster)
bioclim <- stack(list.files(path = "D:/PhD works/world clim rasters/clipped rasters India", pattern = ".tif$", full.names = TRUE))
values_bioclim <- getValues(bioclim)

corr_matrix <- cor(values_bioclim, use = "complete.obs")

library(corrplot)
corrplot(corr_matrix, method = "color", type = "upper", tl.cex = 0.8)

library(caret)
high_corr <- findCorrelation(corr_matrix, cutoff = 0.8)  # set cutoff as needed
bioclim_uncorrelated <- bioclim[[-high_corr]]  # Remove correlated variables
bioclim_uncorrelated


###processing the output and setting decision threshold for the current distribution####

setwd("C:/Users/aritra/Desktop/thrigmopoeinae work/maxent/output") #set path to the folder where the outputs are

library(raster)
library(maps)
library(mapdata)
library(ggplot2)
library(scales)
library(rgdal)

present<-raster("Brachypelma_klassi.asc")
plot(present)
clean_data<-read.csv("cleaned data.csv")
points(clean_data$long,clean_data$lat, pch="+")
extent(present)
extent_to_crop <- extent(70,85,6,20)
cropped_raster <- crop(present, extent_to_crop)
plot(cropped_raster)

##10 percentile training presence##
m <- c(0, 0.082, 0, 0.082, 1, 1)
rclmat <- matrix(m, ncol = 3, byrow = TRUE)
rclmat

rc <- reclassify(present, rclmat)
plot(rc)


##equal training sensitivity and specificity##
m <- c(0, 0.171, 0, 0.171, 1, 1)
rclmat <- matrix(m, ncol = 3, byrow = TRUE)
rclmat

rc <- reclassify(present, rclmat)
plot(rc)


##calculating suitable area
res(rc)

# Calculate the area of each cell in square kilometers
cell_areas <- area(rc)

# Multiply by the binary suitability map
suitable_area_km2 <- sum(cell_areas[values(rc) == 1], na.rm = TRUE)
suitable_area_km2


###processing the output and setting decision threshold for the LGM distribution####

setwd("C:/Users/aritra/Desktop/thrigmopoeinae work/maxent/LGM") #set path to the folder where the outputs are

library(raster)
library(maps)
library(mapdata)
library(ggplot2)
library(scales)
library(rgdal)

present<-raster("Brachypelma_klassi_lgm.asc")
plot(present)
clean_data<-read.csv("cleaned data.csv")
points(clean_data$long,clean_data$lat, pch="+")
extent(present)
extent_to_crop <- extent(70,83,6,20)
cropped_raster <- crop(present, extent_to_crop)
plot(cropped_raster)

##10 percentile training presence##
m <- c(0, 0.082, 0, 0.082, 1, 1)
rclmat <- matrix(m, ncol = 3, byrow = TRUE)
rclmat

rc <- reclassify(present, rclmat)
plot(rc)


##equal training sensitivity and specificity##
m <- c(0, 0.171, 0, 0.171, 1, 1)
rclmat <- matrix(m, ncol = 3, byrow = TRUE)
rclmat

rc <- reclassify(present, rclmat)
plot(rc)


##calculating suitable area
res(rc)

# Calculate the area of each cell in square kilometers
cell_areas <- area(rc)

# Multiply by the binary suitability map
suitable_area_km2 <- sum(cell_areas[values(rc) == 1], na.rm = TRUE)
suitable_area_km2
```
